# Structural Basis of H2B Ubiquitination-Dependent H3K4 Methylation by COMPASS

**DOI:** 10.1101/740738

**Authors:** Peter L. Hsu, Hui Shi, Calvin Leonen, Jianming Kang, Champak Chatterjee, Ning Zheng

**Affiliations:** Department of Pharmacology, Box 357280, University of Washington, Seattle, WA 98195; Howard Hughes Medical Institute, University of Washington, Seattle, WA 98195; Department of Chemistry, University of Washington, Seattle, WA 98195

## Abstract

The COMPASS complex represents the prototype of the SET1/MLL family of methyltransferases that controls gene transcription by H3K4 methylation (H3K4me). Although H2B monoubiquitination (H2Bub) is well-known as a prerequisite histone mark for COMPASS activity, how the H2Bub-H3K4me crosstalk is catalyzed by COMPASS remains unclear. Here, we report the cryo-EM structures of an extended COMPASS catalytic module (CM) bound to the H2Bub and free nucleosome. The COMPASS CM clamps onto the nucleosome disk-face via an extensive interface to capture the flexible H3 N-terminal tail. The interface also sandwiches a critical Set1 arginine-rich motif (ARM) that auto-inhibits COMPASS. Unexpectedly, without enhancing COMPASS-nucleosome interaction, H2Bub activates the enzymatic assembly by packing against Swd1 and alleviating the inhibitory effect of the Set1 ARM upon fastening it to the acidic patch. By unmasking the spatial configuration of the COMPASS-H2Bub-nucleosome assembly, our studies establish the structural framework for understanding the long-studied H2Bub-H3K4me histone modification crosstalk.

## INTRODUCTION

Methylation of histone H3 at lysine 4 (H3K4me) is an evolutionarily conserved post-translational modification that marks actively transcribed genes and regulates transcription in eukaryotic cells. In humans, H3K4 methylation is catalyzed by the SET1/MLL family of histone methyltransferases, which consists of six family members, MLL1–4, SETD1A, and SETD1B (Shilatifard, 2012). These enzymes use the methyl-donor S-adenosylmethionine (SAM) to decorate H3K4 with different degrees of methylation (Ardehali et al., 2011; Denissov et al., 2014; Herz et al., 2012; Hu et al., 2013; Lee et al., 2013; Southall et al., 2009; Wang et al., 2009; Wu et al., 2008).

The catalytic activity of the SET1/MLL enzymes resides in their highly conserved C-terminal SET (<u>S</u>uppressor of variegation, <u>E</u>nhancer of zeste, <u>T</u>rithorax) domains, which are largely inactive on their own (Patel et al., 2009; Shinsky et al., 2014; Southall et al., 2009; Zhang et al., 2015). The SET domains of SET1/MLL methyltransferases must scaffold four binding partners, collectively known as WRAD (<u>W</u>DR5, <u>R</u>BBP5, <u>A</u>SH2L, <u>D</u>PY-30), to form an active catalytic module (CM) that imparts both enzymatic activity and product specificity (H3K4me1, 2 or 3) (Cao et al., 2010; Dou et al., 2006; Patel et al., 2009; Shinsky et al., 2015). Such a requirement is not only conserved in animals, but also in yeast. In fact, yeast Set1, the founding member of the SET1/MLL family, was initially identified and characterized as part of a large multi-protein complex, known as COMPASS (<u>COM</u>plex of <u>P</u>roteins <u>AS</u>sociated with <u>S</u>et1), which features a similar CM (Halbach et al., 2009; Krogan et al., 2002; Mersman et al., 2012; Miller et al., 2001; Nagy et al., 2002; Roguev et al., 2001). By unveiling the architecture of the CMs of COMPASS and COMPASS-like MLL complexes, recent structural studies have shed light on how WRAD activates their SET domains and, together, governs their product specificity (Hsu et al., 2018; Li et al., 2016; Qu et al., 2018). While these studies have illustrated the structural basis of their catalytic activity, how the SET1/MLL enzymatic complexes respond to regulatory signals remains unclear.

Histone H2B monoubiquitination (H2Bub) is a conserved histone mark, which is associated with actively transcribing genes and regulates a myriad of transcriptional processes (Wang et al., 2017). Interestingly, methylation of H3K4 and H3K79, which is catalyzed by COMPASS and DOT1L, respectively, both require H2Bub (Sun and Allis, 2002). Biochemical reconstitution studies have demonstrated the direct activation of these enzymes by the H2B-conjugated ubiquitin (Ub) (Chatterjee et al., 2010; Holt et al., 2015; Kim et al., 2013; McGinty et al., 2008). Despite recent structural studies of H2Bub-dependent H3K79 methylation by DOTL1 (Anderson et al., 2019; Valencia-Sánchez et al., 2019; Worden et al., 2019), how H2Bub activates the multi-subunit COMPASS assembly remains a mystery. Here we report the cryo-electron microscopy (cryo-EM) structures of an extended CM (eCM) of yeast COMPASS bound to the H2Bub modified and unmodified nucleosome. Our results not only reveal how COMPASS recognizes its nucleosomal substrate, but also shed light on how H2Bub allosterically activates the auto-inhibited methyltransferase complex.

## RESULTS AND DISCUSSION

### Reconstitution of H2Bub-dependent COMPASS activity

To gain insight into how the catalytic activity of COMPASS is regulated by H2Bub, we first co-expressed and purified the intact eight-subunit *K. lactis* COMPASS and assembled nucleosome core particles (NCP) containing ubiquitin-modified H2B (uNCPs) (Figure S1A). We and others have previously showed that the five-subunit COMPASS CM complex, which consists of the SET domain of Set1 and the four WRAD subunits, Swd3, Swd1, Bre2, and Sdc1 (Figure 1A), possess a robust activity in methylating H3K4 with unmodified nucleosomes (Hsu et al., 2018; Kim et al., 2013). In contrast, the intact COMPASS exhibited little to no activity against naked nucleosomes (Figure 1B). Remarkably, when presented with ubiquitinated nucleosomes, the full COMPASS complex showed a robust activity of methylating H3K4 to all three methylation states, which is consistent with its H2Bub-dependent activity in the cell. The lack of enzymatic activity in intact COMPASS towards unmodified nucleosomes suggest that the enzymatic complex is somehow inhibited and might not be able to bind the nucleosomal substrate. To our surprise, COMPASS bound nucleosomes almost equally well as the isolated five-subunit catalytic module (Figure 1C and S1B). These results strongly suggest that nucleosome binding by COMPASS is necessary, but not sufficient, for the enzymatic complex to methylate H3K4.

**Figure 1.**
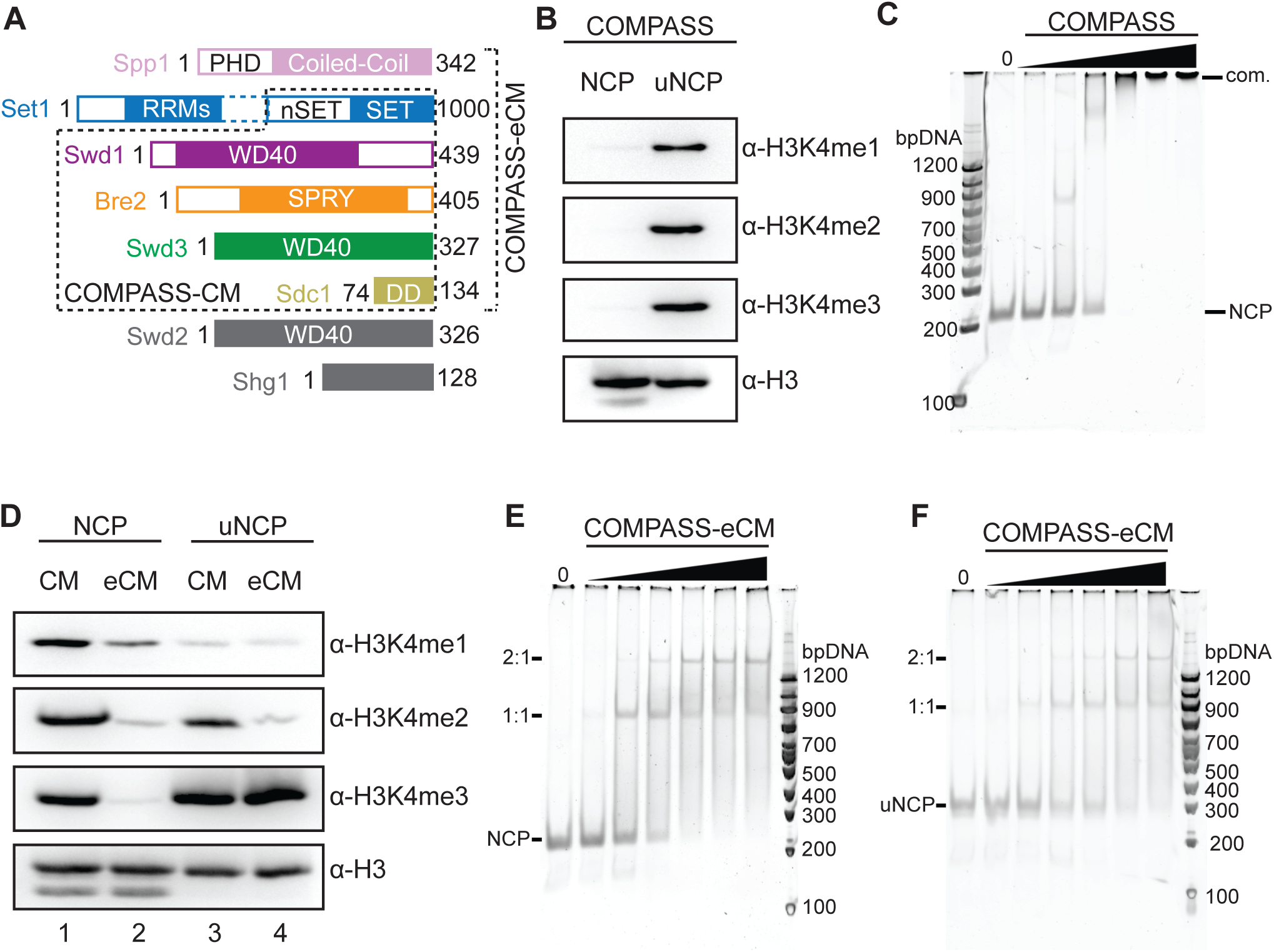
Biochemical analyses of COMPASS nucleosomal binding and activity. A. Domain organization and construct design of COMPASS subunits utilized in biochemical analysis and structure determination. Set1 is not drawn to scale. The dotted box between the nSET and RRM domains indicates a long linker region. The CM boxed in dashed lines contains the Set1 SET domain and the four WRAD subunits. The eCM extends the CM by including the nSET domain and its binding partner, Spp1.
B. H3K4 methyltransferase activity of purified full-length COMPASS on NCP and uNCP substrates. COMPASS exhibits little to no activity on unmodified nucleosomes, while displaying robust methylation on ubiquitinated nucleosomes like its *S. cerevisiae* ortholog.
C. Native TBE gel shift assay of unmodified nucleosomes with increasing concentrations of COMPASS, stained by SybrGold.
D. H3K4 methyltransferase activity of both the COMPASS catalytic module (CM) and extended catalytic module (eCM) on NCP and uNCP substrates. eCM activity is largely suppressed on unmodified nucleosomes and greatly stimulated by H2Bub-nucleosomes.
E. Native TBE gel shift assay of unmodified nucleosomes with increasing concentrations of COMPASS-eCM, stained by SybrGold.
F. Native TBE gel shift assay of H2Bub-nucleosomes with increasing concentrations of COMPASS-eCM, stained by SybrGold.

With a chromatin substrate, an extended six-subunit CM complex (eCM), which includes the nSET domain of Set1 and the Spp1 subunit (Figure 1A), has been previously reported to be largely inactive and represent the minimal core complex with H2Bub-dependent activities (Kim et al., 2013). With the nucleosome substrate, we observed the same stimulating effect of H2Bub on the purified COMPASS eCM (Figure 1D, lane 2 vs. 4). By contrast, the activity of the COMPASS CM was only slightly enhanced by H2Bub (Figure 1D, lane 1 vs. 3). Importantly, the COMPASS eCM shifted nucleosomes just as well as the intact COMPASS and the CM complexes in the gel shift assay (Figure 1C, 1E, and S1B). It also took the same amount of the COMPASS eCM to up-shift NCP in comparison to uNCP (Figure 1F vs. 1E). Together, these results support the notion that the nSET/Spp1 module contains an element that inhibits the intrinsic methyltransferase activity of the CM and confers H2Bub sensitivity to COMPASS. Furthermore, the Ub moiety conjugated to H2B (H2B~Ub) does not increase the substrate-binding affinity of the eCM subcomplex. Instead, it activates the enzyme assembly most likely by reversing the inhibiting effect of the structural element present in the COMPASS eCM outside the CM.

### Overall structure of the COMPASS-uNCP complex

To reveal the structural basis of H2Bub-dependent H3K4 methylation by COMPASS, we prepared a glutaraldehyde cross-linked sample of the COMPASS eCM-uNCP complex and determined its structure by single particle cryo-EM to a global resolution of 3.5 Å (Figure 2 and S2; Table S1). As readily visualized in 2D class averages, the COMPASS eCM can bind to both sides of uNCP, forming a complex at a 2:1 ratio. To obtain a high-resolution map, we performed masked global 3D classification, followed by masked 3D refinement, focusing on the 1:1 enzyme-substrate sub-complex (Figure S2). The final map allowed us to unambiguously dock and rebuild models of the nucleosome and most subunits of the COMPASS eCM assembly (Figure S3). For comparison, we also determined the 3.7 Å resolution cryo-EM structure of the COMPASS eCM bound to an unmodified nucleosome. This Ub-free structure reveals a similar overall architecture with discernable local differences (e.g. Figure S3C vs. S3D), which will be highlighted after the description of the COMPASS eCM-uNCP structure.

**Figure 2.**
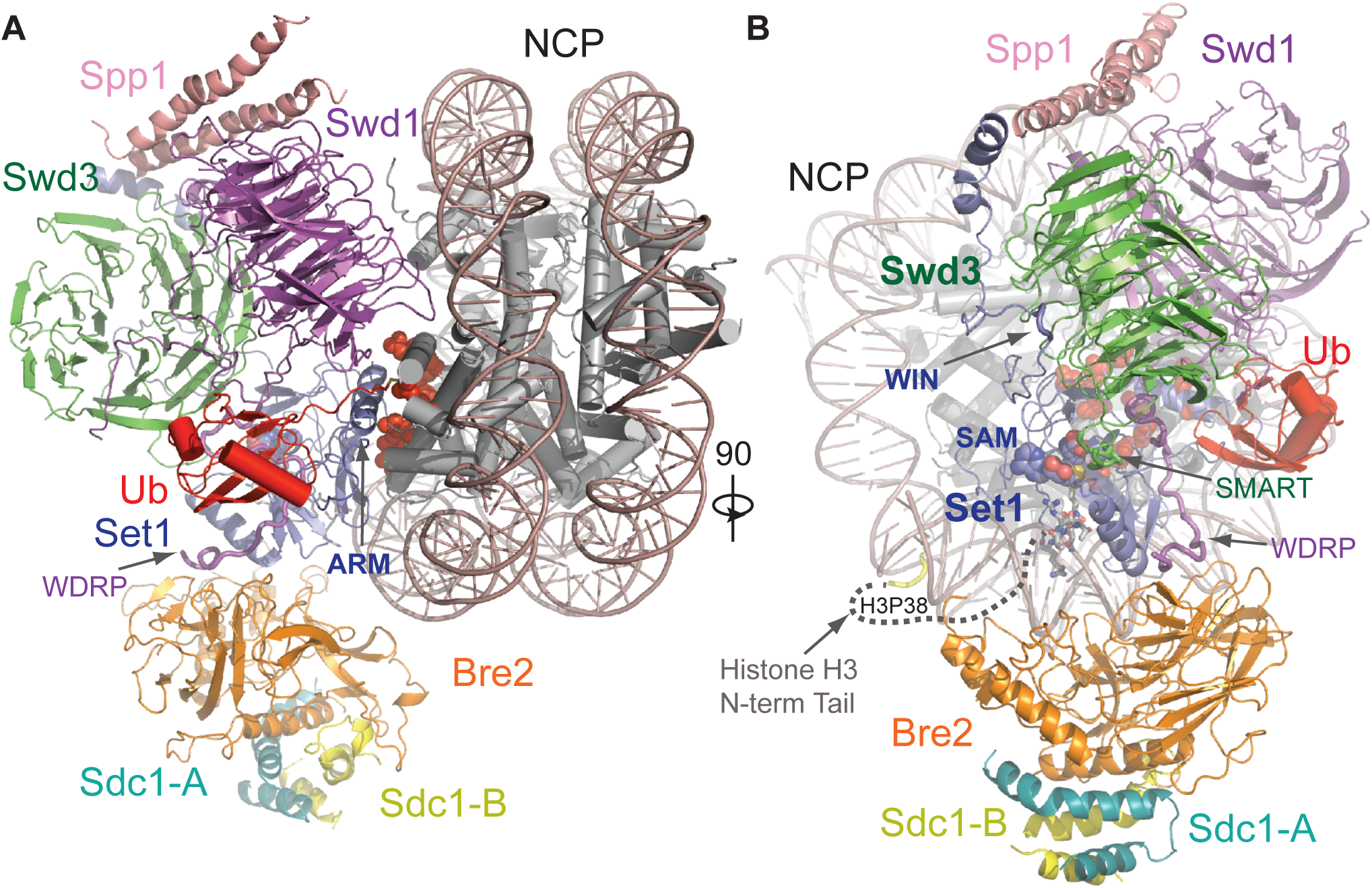
Overall structure of the COMPASS eCM-uNCP complex. A. Model of COMPASS-eCM bound to the H2Bub modified nucleosome. Set1 (blue), Swd1 (magenta), Swd3 (green), Bre2 (orange), Sdc1-A and B (turquoise and yellow), Spp1 (pink), and Ub (red) are shown in cartoon form. Nucleosomal DNA (dark salmon) is shown in cartoon. The histone octamer is shown in cylinders. The SAM cofactor is shown in space-filling form. Residues forming the H2A-H2B acidic patch are shown in red spheres. The positions of the Swd1 WDRP and Set1 ARM helix are indicated by arrows.
B. View of (A) rotated 90° and colored in the same scheme. The positions of the Set1 WIN motif (tube form), Swd3 SMART motif (tube), and Swd1 WDRP (tube) are indicated by arrows. A dashed line indicates the flexible histone H3 N-terminal tail spanning residue 9 to 38, which connects the first stretch of H3 seen in the NCP body (yellow tube) and the H3 (A1-R8) peptide (sticks) found in the Set1 SET domain.

The COMPASS eCM recognizes two distinct parts of its nucleosomal substrate, the surface of the well-ordered NCP disk body and the distal end of the flexible histone H3 N-terminal tail. Upon binding to the nucleosome, the enzymatic complex partitions H2B~Ub and the histone H3 N-terminal tail on its two separate sides (Figure 2B). The COMPASS eCM spans across the entire disk face of the nucleosome and uses four out of six subunits to make direct contacts with all histones and nucleosomal DNA (Figure S4A). Only Swd3 and Sdc1 are isolated from the interface, in support of their roles as structural subunits of the COMPASS assembly. The COMPASS eCM buries a total of ~3000 Å^2^ surface area on the nucleosome, consistent with the strong association between the two complexes.

Similar to its central location in the isolated COMPASS eCM, the Set1 SET domain is positioned at the center of the COMPASS eCM-uNCP interface in close proximity to Ub (Figure 2). We were able to observe clear densities for both the SAM cofactor, as well as the entire tip of the H3 N-terminal tail (amino acid 1 to 8) at the active site of the SET domain (Figure S3C), suggesting that the COMPASS eCM was crosslinked and captured in its pre-reaction state. The remainder of the H3 tail ranging from amino acid 9 to 37 is invisible in our maps, likely due to its intrinsic flexibility. The H3K4 peptide can only weakly bind to the SET1/MLL enzymatic complex with an affinity of ~100 μM (Zhang et al., 2015). The overall bipartite binding mode of the COMPASS eCM to the nucleosome not only underlines the ability of the Set1 catalytic domain to differentiate H3K4 from other histone lysine residues, but also accentuate the contribution made by the interactions between COMPASS and nucleosome disk body to target site engagement.

Upon conjugated to H2B, Ub uses its canonical Ile44 hydrophobic patch as well as its C-terminal tail to pack against Swd1. Meanwhile, H2B~Ub also interacts with a long Set1 α-helix adopted by an arginine-rich motif (ARM) immediately preceding and outside the SET domain in sequence (Figure 2A). Unlike many other chromatin-binding proteins (Figure S4) (McGinty and Tan, 2016), the COMPASS CM subunits shared among all SET1/MLL enzymes do not interact with, but instead arches over, the H2A-H2B acidic patch (Figure 2A). Interestingly, the space in between is occupied by the Set1 ARM helix, which is evolutionarily conserved but unique to fungi Set1 proteins and their mammalian orthologs, SETD1A and SETD1B, whose enzymatic activities are H2Bub-dependent. The Set1 ARM helix stands out as the hallmark of the COMPASS eCM-uNCP assembly by not only anchoring its N-terminal half right next to the SET domain-nucleosome interface, but also employing its C-terminal half to directly contact H2B~Ub.

### Bre2 and Spp1 bind nucleosomal DNA

Previous studies have suggested that protein-nucleic acid interactions are critical for COMPASS and COMPASS-like complexes to recognize and methylate substrates (Chen et al., 2011; Trésaugues et al., 2006). The COMPASS eCM-uNCP complex structure reveals multiple DNA contacts made by Spp1 (CFP1 in humans) and the WRAD subunit Bre2 at the two extreme ends of the interface (Figure 3A). Although clear densities were observed for the majority of the nucleosome-COMPASS interface, the local resolution for Spp1 and Bre2-Sdc1, which are at the periphery of the complex, was not high enough for manual model building (Figure S3A). Instead, we fit a homology model of the central coiled-coil region of *K. lactis* Spp1 derived from the structure of its *S. cerevisiae* ortholog to the map (Qu et al., 2018). We observed discernable densities contacting the major groove of nucleosomal DNA that corresponds to a region missing in *S. cerevisiae* Spp1 (Figure 3B). This Spp1 region sits at the junction between the two long α-helices and contains a series of positively charged residues that is mostly disordered in the absence of the nucleosomal substrate, but likely involved in engaging DNA (Figure 3C).

**Figure 3.**
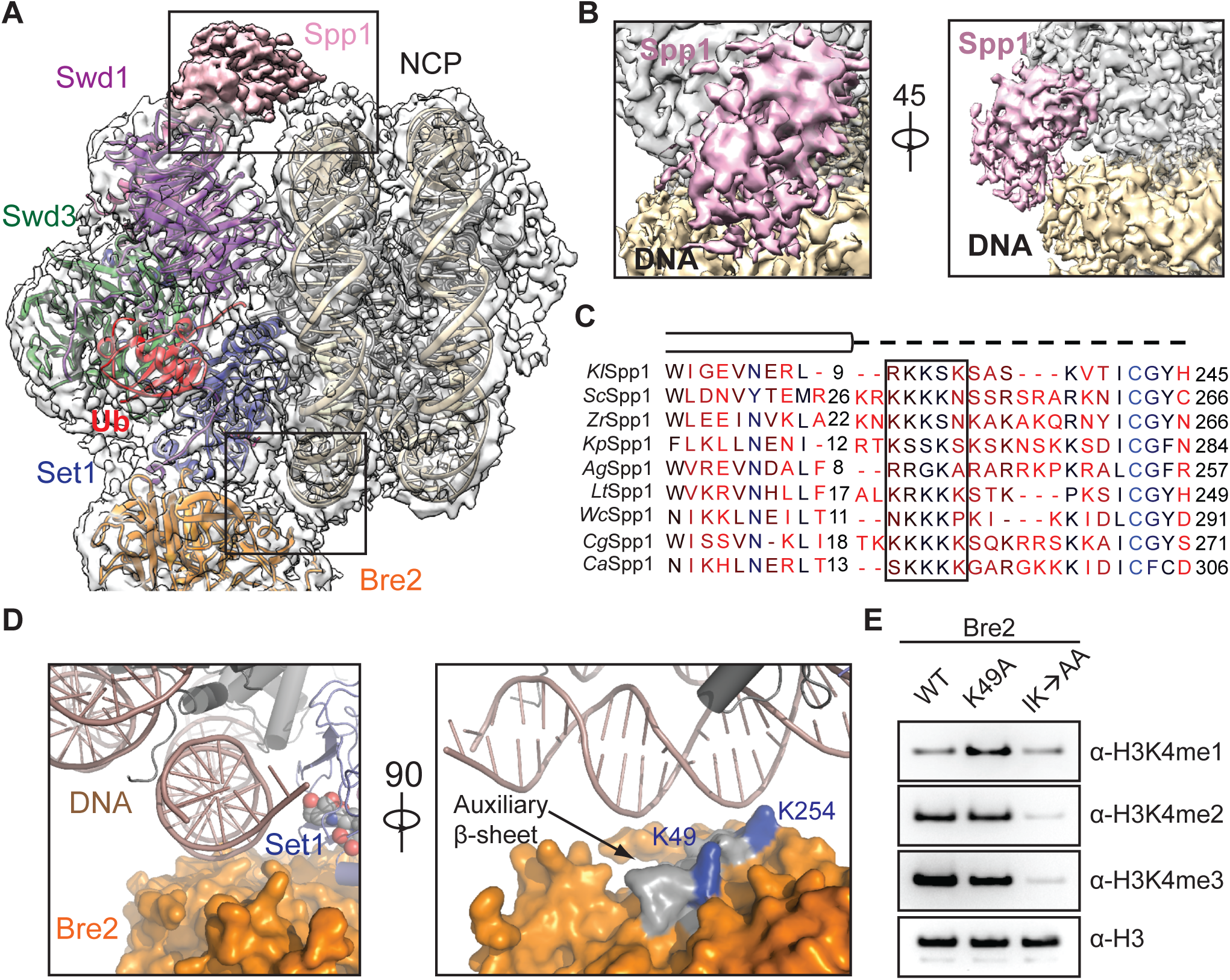
DNA binding stabilizes COMPASS on nucleosomes. A. Zoom out view of the complex structure superimposed on the cryo-EM map. Major DNA binding sites are boxed. Swd1 is shown in magenta, Swd3 in green, Set1 in blue, Bre2 in orange, Spp1 in pink, Ub in red, histones in dark grey, and nucleosomal DNA in dark salmon.
B. Close up views of the Spp1 DNA binding domain density (pink) on nucleosomal DNA (wheat) in two different orientations.
C. Structure-based sequence alignment of yeast Spp1 orthologs from *K. lactis* (*Kl*), *S. cerevisiae* (*Sc*), *Z. rouxii* (*Zr*), *K. pastoris* (*Kp*), *A. gossypii* (*Ag*), *L. thermotolerans* (*Lt*), *W. ciferri* (*Wc*), *C. glabrata* (*Cg*), and *C. albicans* (*Ca*). Cylinders denote α-helices, while the dashed line indicates an unmodeled region, predicted to lack secondary structure. A basic region with potential DNA binding function is boxed.
D. (Left panel) Close up view of Bre2 (orange in surface representation) looking down the nucleosomal DNA entry/exit point (cartoon representation, dark salmon). (Right panel) 90° rotation of the left panel, highlighting the position of auxiliary β-sheet (grey) on Bre2 relative to DNA. Two highly conserved lysine residues on the surface of this structural element are colored in blue.
E. H3K4 methyltransferase activity of the COMPASS catalytic module (CM) assembled with Bre2 mutants against nucleosomal substrates. The IK→AA double mutant of two conserved surface residues, Ile251 and Lys254, drastically reduces activity of the CM against H3K4.

Bre2 mirrors the action of Spp1 by interacting with DNA at the opposite end of the nucleosome (Figure 3A). As previously revealed, Bre2 contains a non-canonical SPRY domain, which is characterized by a twisted auxiliary β-sheet harboring a cluster of highly conserved but solvent-exposed residues (Hsu et al., 2018). At the COMPASS-DNA interface, Bre2 embraces the phosphate backbone of nucleosomal DNA using the edge of its auxiliary β-sheet (Figure 3D). To assess the significance of this interface, we purified COMPASS CM with two sets of Bre2 mutations, focusing on two highly conserved lysine residues on the surface of the auxiliary β-sheet that are in close proximity to the phosphate backbone of DNA (Figure 3D, right panel). Compared to the wild type protein, mutation of Bre2 Lys49 showed a minor effect on the methyltransferase activity, whereas a Bre2 double mutant, Ile251Ala and Lys254Ala (IK→AA), severely compromised the production of H3K4 in all three methylation states (Figure 3E). As expected, this loss of enzymatic activity is attributable to the disruption of nucleosome binding by the dual mutation as assessed by gel shift assays (Figure S1B and C). Taken together, these data suggest that DNA binding by Bre2 is critical for the recruitment and activity of COMPASS on nucleosomes.

### Swd1 contacts all four histones and nucleosomal DNA

Swd1 plays a critical role in organizing the COMPASS eCM by interacting with almost every subunit except Sdc1 (Halbach et al., 2009; Hsu et al., 2018; Qu et al., 2018). This functional importance of Swd1 is further magnified by the COMPASS-uNCP structure, in which Swd1 closely interacts with all four core histones, nucleosomal DNA, and Ub. Using the edge of its β-propeller domain and several loop regions, Swd1 docks to the nucleosome disk at a pronounced three-helix cleft formed by the last two helices of H2B (α3 and αC), the second helix of H2A (α2), and DNA (Figure 4A and 4B). As a canonical WD40 repeat-containing protein (Sprague et al., 2000), Swd1 utilizes the conserved loop connecting the “D” strand of blade 5 and the “A” strand of blade 6 (5D-6A loop), as well as the solvent exposed “D” strand of blade 6 to cradle the long C-terminal αC helix of H2B (Figure 4B and 4C). Meanwhile, the tip of the Swd1 5D-6A loop is deeply inserted into the three-helix cleft. Two consecutive Swd1 isoleucine residues, Ile271 and Ile272, interact with a hydrophobic patch formed around H2B Val115 and H2A Tyr50. The interface is enhanced by two nearby invariant Swd1 residues, Asn273 and Arg274, which make a hydrogen bond with H2B Gln96 and a salt bridge with the DNA backbone, respectively (Figure 4D and 4E).

**Figure 4.**
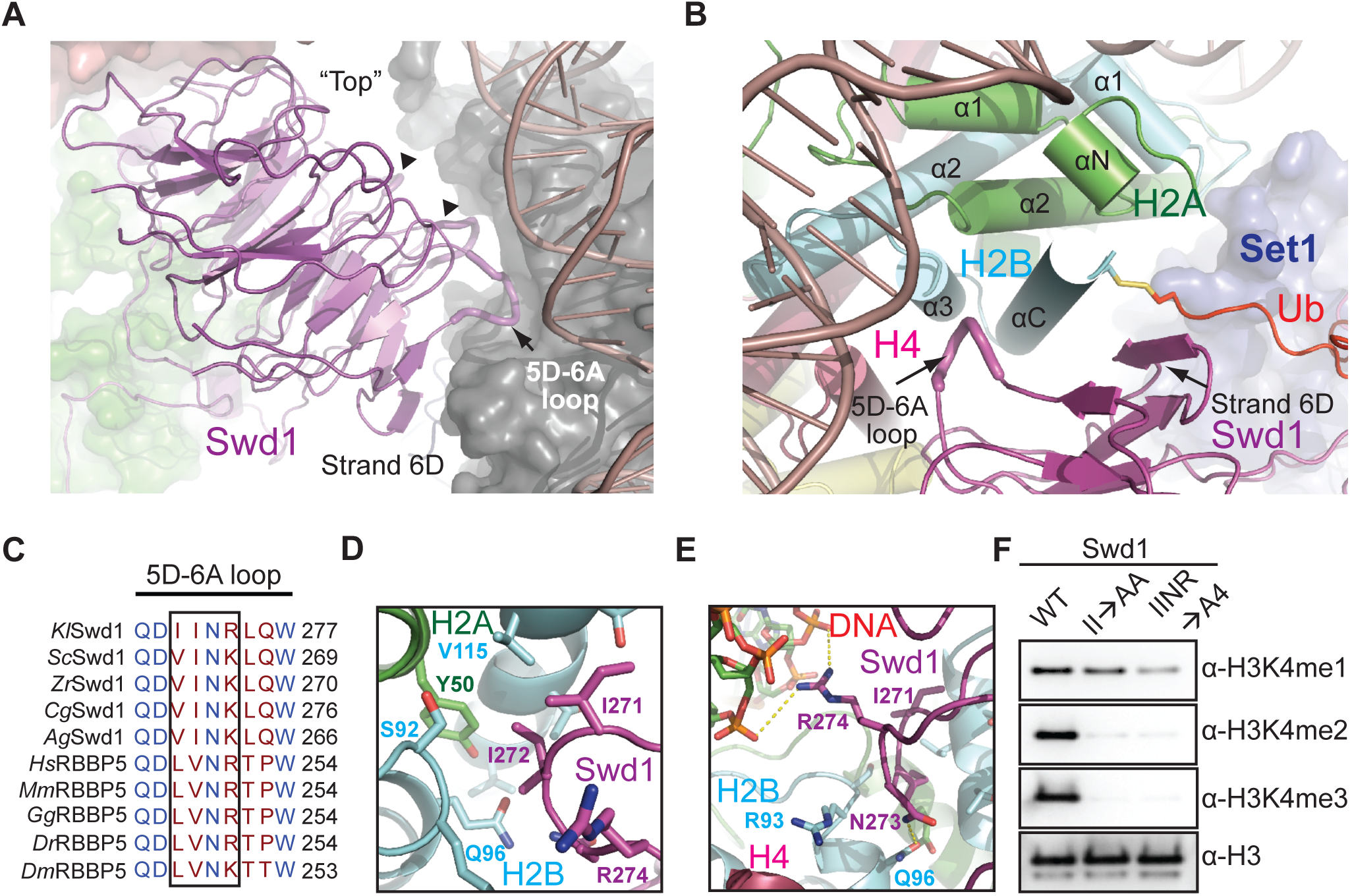
Swd1 is a histone binding protein. A. Side view of Swd1 (magenta, cartoon) positioned over the histone surface (dark grey, surface). The “top” face of the WD40 is pointed towards the nucleosome, while the “bottom” is facing COMPASS, interacting with Swd3 (green, surface). The “top” surface loops interacting with the histone surface are shown in tube representation. The 5D-6A loop (tube form) and strand D from blade 6 are labeled. The two loops interacting with H3 and H4 are indicated by triangles.
B. Top down view, looking down the C-terminal H2B helix (cyan). Swd1 (magenta) wraps around this helix using the 5D-6A loop (tube form) and strand 6D. Set1 is shown in blue surface representation. H2A and H4 are shown in green and deep red respectively in cartoon form. The C-terminal tail of Ub (red) is fused to H2B via a disulfide linkage.
C. Local structure-based sequence alignment of the 5D-6A loop across *K. lactis* (*Kl*), *S. cerevisiae* (*Sc*), *Z. rouxii* (*Zr*), *C. glabrata* (*Cg*), *A. gossypii* (*Ag*), *H. sapiens* (*Hs*), *M. musculus* (*Mm*), *G. gallus* (*Gg*), *D. rerio* (*Dr*), and *D. melanogaster* (*Dm*) orthologs. Straight line denotes an element with no secondary structure. The 4-residue segment that inserts itself into the H2A-H2B 3-helix cleft is boxed.
D. Close up view centered on the two conserved hydrophobic residues in the 5D-6A loop (magenta) at the 3-helix juncture (H2B in cyan, H2A in green).
E. Close up view centered on the two polar residues in the 5D-6A loop (magenta). Asn273 forms a hydrogen bond with H2B Gln96 (cyan), while Arg274 interacts with the phosphate backbone of nucleosomal DNA (red, sticks). Polar and salt bridge interactions are shown as dotted yellow lines.
F. H3K4 methyltransferase activity of the COMPASS catalytic module (CM) assembled with Swd1 5D-6A loop mutants against nucleosomal substrates. A double mutant of two hydrophobic residues (II→AA) reduces methyltransferase activity to mono-methylation. A quadruple mutant (IINR→A4) nearly renders the CM inactive.

The importance of the conserved 5D-6A loop of Swd1 in locking COMPASS to nucleosomes is underscored by alanine substitutions in this loop. Mutation of its two central hydrophobic residues, Ile271A/Ile272A (II→AA), effectively abolishes H3K4 di- and tri-methylation by the COMPASS CM (Figure 4F). Alanine replacement of all four residues in the loop (IINR→A4) reduces activity of the COMPASS CM to near background levels for all three methylation states. Importantly, the tetra-mutant significantly weakens the binding of the COMPASS CM to nucleosomes, indicating that Swd1 is on par with Bre2 in promoting nucleosomal substrate recognition by the histone methyltransferase complex (Figure S1D).

### Binding of the Set1 catalytic domain to H2A

The Set1 catalytic domain represents the heart of the COMPASS CM. It is held by a highly conserved WD40 repeat proximal (WDRP) region of Swd1 and is sandwiched between Bre2 and the two CM β-propeller subunits, Swd1 and Swd3 (Figure 2). Superposition analysis suggests that we have captured the SET domain in a closed conformation that is conducive for catalysis (Figure S5A). In the COMPASS eCM-uNCP structure, the Set1 SET domain is placed at the center of the enzyme-substrate assembly, directly packing against the α2 helix of H2A through a marked interface (Figure 5A). Upon engaging the nucleosome, the N-terminal α-helix of the SET domain unravels into a more linear configuration (Figure 5B and S5B), which enables Lys855 and Arg856 to interact with three H2A residues: Glu64, Asn68 and Asp72, on its α2 helix (Figure 5C). The Set1-H2A interaction is further bolstered by a quintet of hydrophobic Set1 residues, Leu853, Met883, Ile948, Val950, and Val957, which are presented by the β-sheet of the SET-N/C subdomain. Together, these residues wrap around the C-terminal tip of the H2A α2 helix and bury H2A Arg71 that is neutralized by the nearby H2B Asp52 residue (Figure 5D). Noticeably, these hydrophobic residues are not only conserved across all Set1 orthologs, but also found in all the SET domains of the expanded SET1/MLL family in mammals (Figure S5C). The binding mode of the catalytic SET domain to H2A, therefore, might be common to all SET1/MLL enzymatic complexes.

**Figure 5.**
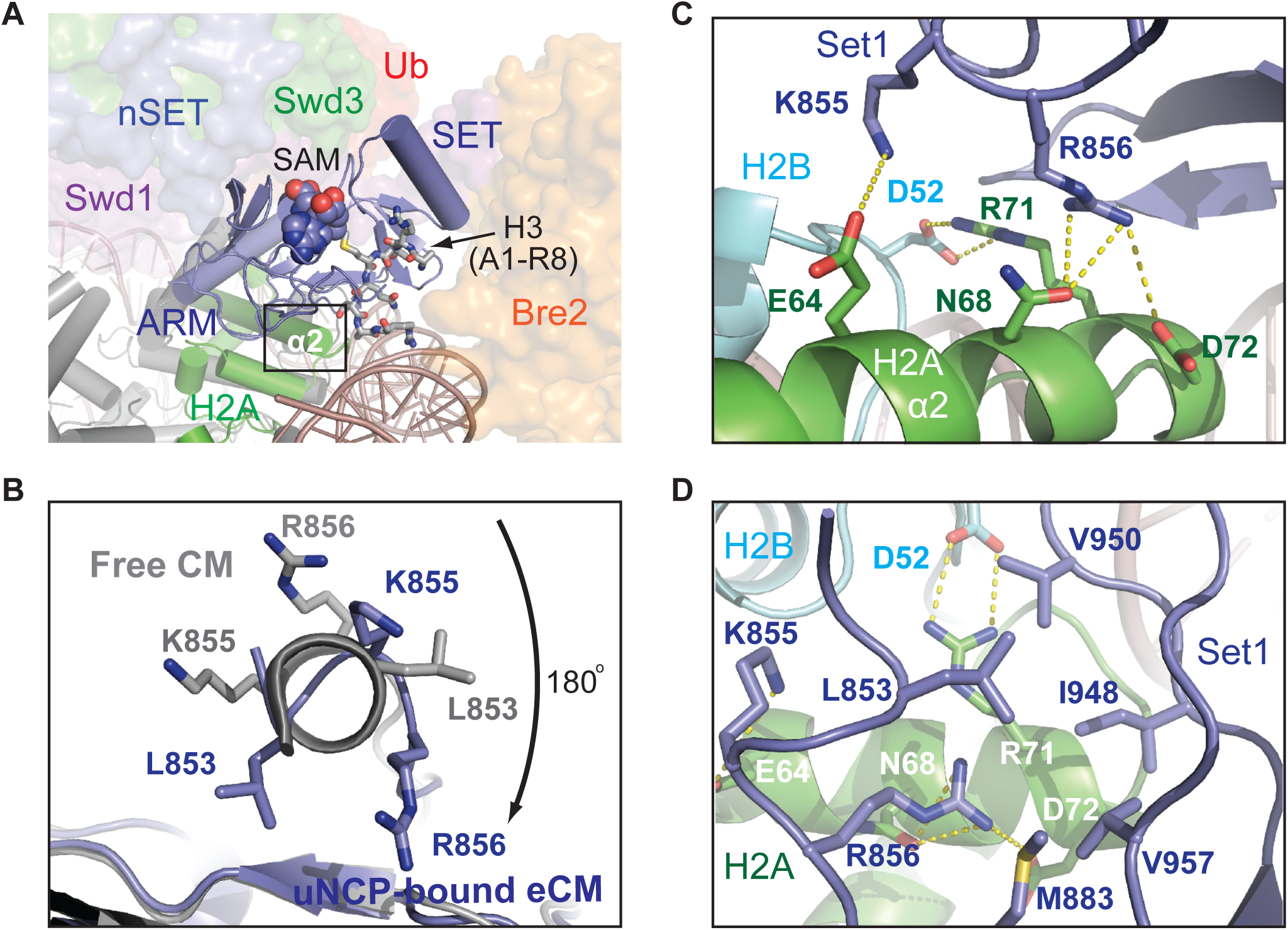
The SET N-terminus unravels to bind H2A. A. Zoom out view of Set1-SET (blue, cartoon) being positioned over histone H2A (green) and sandwiched by Bre2 (orange, surface) and Swd1 (magenta, surface). Swd3 (green) and the nSET domain (blue) are also shown in surface representation. The cofactor SAM and the H3 N-terminus are shown in space-filling and stick form, respectively. Boxed is the region of interaction between Set1 and H2A.
B. View looking down the N-terminal helix of the Set1 SET domain. The SET domain found in the NCP-bound structure is shown in blue, while the free form of the SET domain is shown in grey. Residues involved in interacting with H2A upon helical unraveling are shown in sticks.
C. Close up view of the rearranged Set1 N-terminus (blue) presenting both Lys855 and Arg856 into a polar tip found on H2A α2 helix (green). Hydrogen bonds and salt bridges are shown as dashed yellow lines.
D. Close up view of the Set1 hydrophobic claw (blue) interaction with H2A (green). H2B is shown in cyan. H3 and H4 are omitted for clarity. Set1, H2A, and H2B amino acids are labeled in blue, white, and cyan, respectively.

A previous systematic alanine scanning analysis of all four core histones in yeast has identified four histone H2A residues, whose mutations impaired H3K4 di-and trimethylation *in vivo* (Nakanishi et al., 2008). Strikingly, three of these residues are localized at the SET domain-H2A interface and perfectly match to the three H2A α2 helix amino acids, Glu64, Asn68, and Asp72, that directly interact with the Set1 SET domain (Figure 5C). Proper packing of the Set1 catalytic domain against histone H2A, thus, is critical for COMPASS to maintain its catalytic competency in the context of the nucleosome.

### The Ubiquitin-COMPASS interface

The H2B-conjugated Ub moiety makes close interactions with the nucleosome-bound COMPASS eCM via multiple interfaces. Its β-grasp fold not only snugly fits into a concave surface presented by several spatially conglomerated Swd1 structural elements, but also packs against a platform formed by the Set1 sequence immediately preceding the catalytic domain (Figure 2 and 6A). As seen in many Ub-protein interfaces (Winget and Mayor, 2010), H2B~Ub uses its Ile44-Val70-Leu8 hydrophobic patch to adhere to a hydrophobic surface formed among Phe10 and Leu13 of the Swd1 N-terminal extension and Val409 of the Swd1 C-terminal tail (Figure 6B). This hydrophobic interface is stabilized on both sides by two nearby positively charged Ub residues, Lys48 and Arg42, which form a salt bridge with Glu15 and Glu405 of Swd1, respectively. The H2B~Ub-Swd1 interface is further extended into the Ub tail, whose two arginine residues, Arg72 and Arg74, cling to the Swd1 β-propeller via additional polar interactions. Overall, a continuous surface area of H2B~Ub that spans about 30 Å is exclusively recognized by Swd1 at this interface, suggesting that the same interactions could take place between H2B~Ub and other SET1/MLL enzymatic complexes.

**Figure 6.**
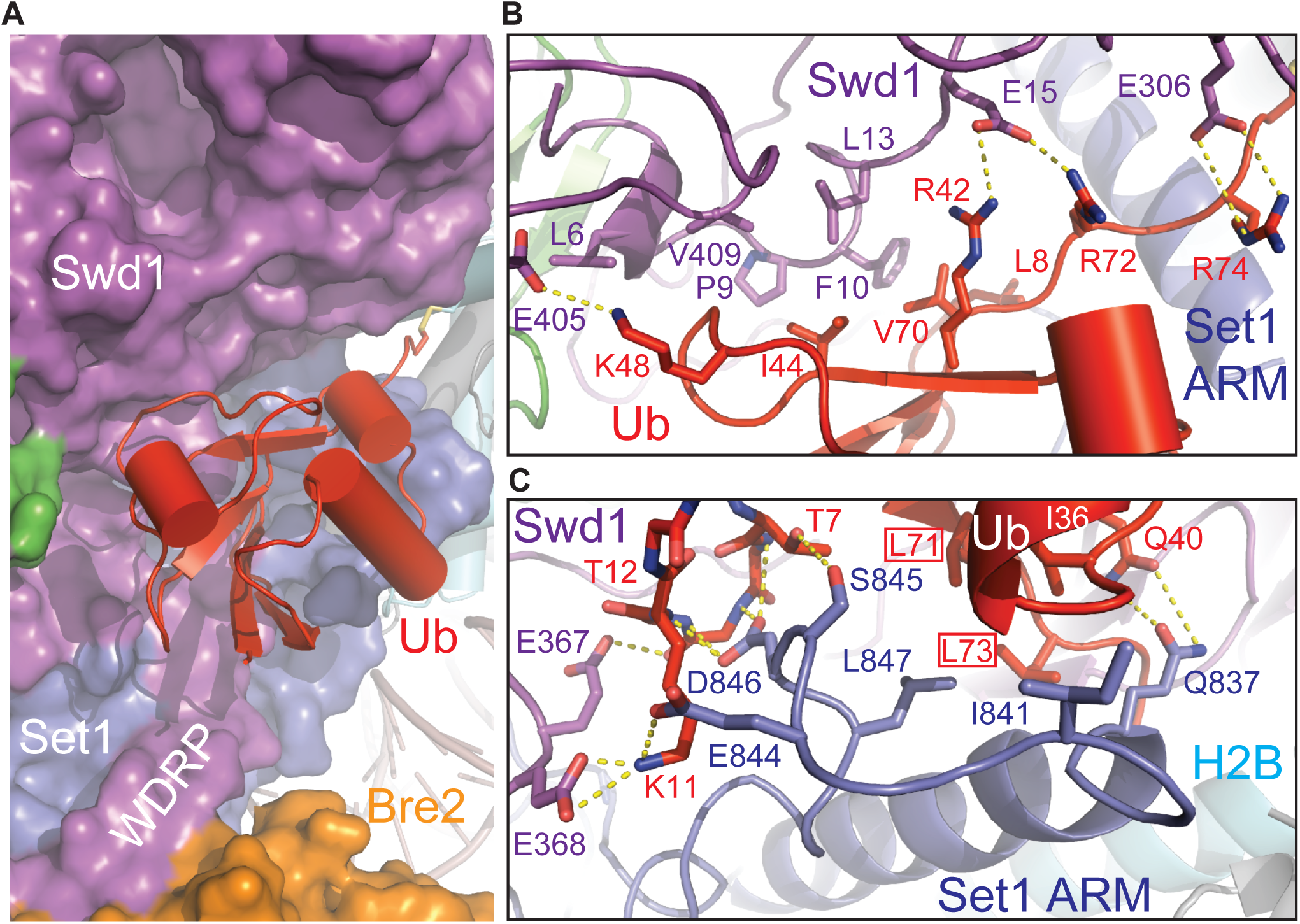
H2B~Ub packs against Swd1 and Set1. A. Zoom-out view of H2B~Ub (red, cartoon) against COMPASS. Swd1 (magenta), Set1 (blue), and Bre2 (orange) are shown in surface form. The disulfide linkage between H2B (cyan, cylinder) and Ub is shown in sticks. The critical WDRP necessary for Set1 activity is labeled.
B. Close up view of the interface between the Ub Ile44 hydrophobic patch and Swd1. Ub packs extensively against a series of conserved residues on Swd1 spread across its N-terminus, WD40, and C-terminal tail. Key side chains are shown in sticks and labeled. Hydrogen bonds and salt bridges are shown as dashed yellow lines.
C. Close up view of the interface between the Set1 SET N-terminus (blue) and Ub Ile36 underside site (red). The N-terminus of the SET domain makes several backbone interactions with Ub between Thr7 and Thr12 (shown in sticks, side chains omitted for clarity). A series of Set1 hydrophobic residues pack against the base of the Ub helix. The Ub Leu71/Leu73 pair identified in Holt et al., 2015 critical for H2Bub-H3K4me crosstalk are boxed. Hydrogen bonds and salt bridges are shown as dashed yellow lines.

In addition to the Ile44-centered hydrophobic patch, H2B~Ub also engages with COMPASS via its Ile36 site situated at a separate location of the β-grasp fold (Winget and Mayor, 2010). Upon unraveling, the N-terminal α-helix of the SET domain and its preceding sequence adopt a highly coiled structure, which inserts a loop into the H2B~Ub pocket demarcated by Ile36, the β1-β2 loop, and the Ub tail (Figure 6C). Within this pocket, Leu847 of Set1 packs against Leu71 and Ile36 of H2B~Ub, while its three preceding residues, Glu844, Ser845, and Asp846, make multiple polar interactions with the β1-β2 loop of Ub on the opposite side. Moreover, the Swd1 WDRP loop joins in at this interface by stabilizing the Ub Lys11 residue with two negatively charged residues. Strikingly, both Leu71 and Leu73 have been previously identified by an alanine scan of Ub as essential for H2Bub-dependent H3K4 methylation (Holt et al., 2015), indicating that the interface formed between Ub and the unraveled SET N-terminus is critical for productive catalytic activation. Furthermore, the majority of the Set1 residues at this three-molecule junction are variable in mammalian Set1 paralogues, MLL1–4, suggesting that this Set1-H2B~Ub interface is most likely unique to fungi Set1 and its mammalian SETD1A and SETD1B orthologues. In fact, this Set1-specific property is manifested by the further upstream ARM helix of the methyltransferase, which is only found in Set1 orthologues.

### Docking of the Set1 ARM helix to acidic patch

Previous studies have identified an arginine-rich motif flanked by the Set1 nSET and SET domains that plays a critical role in mediating H2Bub-dependent H3K4 methylation by the COMPASS eCM (Kim et al., 2013) (Figure 7A). This Set1 motif contains three invariant arginine residues conserved from yeast to humans. Although the Set1 ARM is completely disordered in the free COMPASS eCM structure (Qu et al., 2018), it becomes buried at the COMPASS-uNCP interface and folds into a long α-helix that is directly connected to N-terminus of the SET domain.

**Figure 7.**
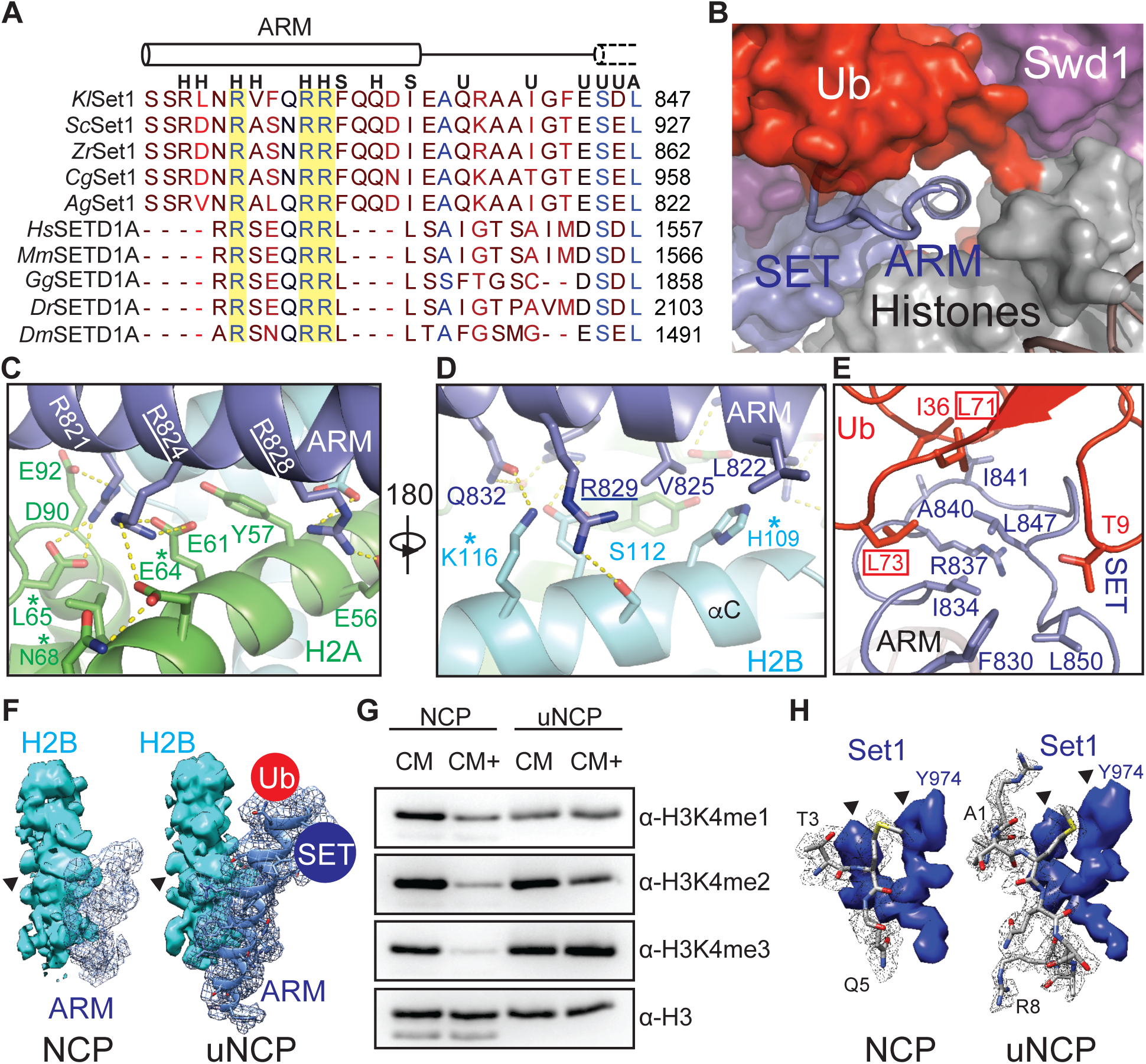
The Set1 ARM regulates COMPASS H2Bub sensitivity. A. Sequence alignment of the nSET Arg-rich motif across *K. lactis* (*Kl*), *S. cerevisiae* (*Sc*), *Z. rouxii* (*Zr*), *C. glabrata* (*Cg*), *A. gossypii* (*Ag*), *H. sapiens* (*Hs*), *M. musculus* (*Mm*), *G. gallus* (*Gg*), *D. rerio* (*Dr*), and *D. melanogaster* (*Dm*) orthologs. The dotted cylinder represents the unraveled SET N-terminal helix. Cylinders represents helices, and lines indicate coiled structure. The strictly conserved arginine residues are highlighted by a yellow background. Letters above the alignment indicate the interactions the residue is involved in. **H** indicates a histone interaction, **S** indicates a SET domain interaction, **U** indicates a Ub interface, and **A** indicates a Set1 ARM interaction.
B. View of the tunnel formed by the junction between Set1 (blue, surface), Swd1 (magenta, surface), Ub (red, surface), and the histone octamer (grey, surface). Swd3 sits at the periphery (green, surface). The Set1 ARM helix (blue, cartoon) sits in the center of this tunnel. The loop connecting the SET domain and the ARM helix is shown in tube form.
C. Close up view of the Set1 ARM helix (blue) against the acidic surface of H2A (green). Side chains are shown in sticks. Two of the three strictly conserved arginine residues that impair H2Bub-H3K4me crosstalk (Kim et al., 2013) are underlined. H2A residues identified in (Nakanishi et al., 2008) critical for H2Bub dependent H3K4 methylation are marked by an asterisk. Hydrogen bonds and salt bridges are shown as dashed yellow lines.
D. Close up view of the Set1 ARM (blue) against the H2B αC helix (cyan). Side chains are shown in sticks. The third strictly conserved arginine residue involved in the H2Bub-H3K4me crosstalk (Kim et al., 2013) is underlined. H2B residues identified in Nakanishi et al., 2008 important for H2Bub dependent H3K4 methylation are marked by an asterisk. Hydrogen bonds and salt bridges are shown as dashed yellow lines.
E. Close up view of the three-way interface between the Set1 (blue) ARM, SET N-terminus, and the Ub (red) Ile36 site. Side chains are shown in sticks. The ARM helix packs against the SET N-terminus using an interlocking series of hydrophobic amino acids. Ub contributes to this interface by helping forming the connecting loop between the ARM and SET regions through hydrophobic packing. Ub Leu71/Leu73 identified by (Holt et al., 2015) to be important for H2Bub stimulated H3K4me activity are boxed.
F. Comparison of the Set1 ARM density (blue mesh) between the unmodified NCP-bound (left) and uNCP-bound (right) structure of the COMPASS eCM. The neighboring H2B density (cyan, surface) was used to normalize the ARM densities. The key side chain feature on H2B used for controlling the map contour level is indicated by a triangle. The uNCP density has the model of the ARM superposed in the mesh. The positions of where Ub and the SET domain dock on the ARM helix are indicated by a red and blue circle, respectively.
G. H3K4 methyltransferase activity of both the COMPASS catalytic module (CM) and catalytic module extended (CM+) on NCP and uNCP substrates. CM+ activity is suppressed relative to the CM on NCPs, and regains robust methyltransferase activity on uNCPs.
H. Comparison of the H3 N-terminal tail density (grey mesh) at the Set1 catalytic cleft between the unmodified NCP-bound (left) and uNCP-bound (right) structure of the COMPASS eCM. The density of a nearby Set1 sequence was used to normalize the H3 N-terminal tail densities. The key features of the Set1 density for controlling the map contour level is indicated by triangles. The models of the H3 N-terminal tail centered at the H3K4 methylation site are shown in sticks.

In the COMPASS-uNCP complex, the Set1 ARM helix runs through the tunnel formed between the COMPASS eCM and the nucleosome disk body and is firmly locked down to the nucleosomal acidic patch (Figure 7B) (McGinty and Tan, 2016). At the bottom side of the Set1 ARM helix, two out of the three strictly conserved arginine residues, Arg824 and Arg828, each make multi-valent polar interactions with a distinct pair of negatively charged histone acidic patch residues (Figure 7C). They are joined by an upstream arginine residue, Arg821, and four H2A residues, Tyr57, Leu65, Asp90, and Glu92, which further augment the complementary interface. Next to the acidic patch, the Set1 ARM helix is held by the H2B αC helix, which employs two polar residues, Ser112 and Lys116, to form a hydrogen bond with Arg829 and Gln832 of Set1, respectively (Figure 7D). The Set1 ARM helix is further secured by its two hydrophobic residues, Leu822 and Val825, which interact with His109 of H2A through van der Waals packing.

A subset of the H2A-H2B residues interacting with the Set1 ARM helix have also been previously identified in the systematic alanine scanning analysis of histone residues that are required for proper H2Bub-mediated H3K4 methylation (Nakanishi et al., 2008). Specifically, alanine substitution of the H2B resides equivalent to His109 and Lys116, as well as Glu64 and Leu65 of H2A all negatively impacted H3K4 di- and trimethylation *in vivo* (Figure 7C, D). These previously published results not only validate the specific Set1 ARM-H2A-H2B interface revealed in the COMPASS eCM-uNCP structure, but also echo the loss of function of the Set1 ARM in supporting H2Bub-dependent activity of the COMPASS eCM due to Arg→Ala mutations (Kim et al., 2013).

### Effects of H2B~Ub on Set1

Upon anchoring at the nucleosomal acidic patch, the Set1 ARM helix also interfaces with and, thereby, bridges two central components of the COMPASS eCM-uNCP complex, the Set1 catalytic SET domain and H2B~Ub. At one end, the N-terminal half of the Set1 ARM helix buttresses the Set1 catalytic domain that rests on the H2A α2 helix (Figure 5A and 7B). At the other end, the C-terminal half of the same helix presents two hydrophobic and an arginine residue, Phe830, Ile834, and Arg837, to hold onto the highly coiled N-terminal sequence of the SET domain. Together, they give rise to a platform that binds to both the Ile36 site and the C-terminal tail of Ub (Figure 7E).

The central and strategic location of the Set1 ARM helix in the COMPASS eCM-uNCP structure and its crucial role in mediating H2Bub-dependent H3K4 methylation by COMPASS strongly suggests that it is a key factor that sensitizes COMPASS to H2Bub. To better understand the nature the H2Bub-H3K4me crosstalk, we determined the cryo-EM structure of the COMPASS eCM bound to the unmodified NCP (Figure S6) and compared it with the Ub-conjugated form of the nucleosome. In the absence of H2B~Ub, the COMPASS eCM docks to the nucleosome in an overall similar manner as it does to uNCP (Figure S7A). However, the density corresponding to the C-terminal half of the Set1 ARM helix and the N-terminal region of the SET domain is entirely missing (Figure 7F). By contrast, we were able to locate the density for the N-terminal half of the helix, which occupies the acidic patch. This result corroborates the idea that the Set1 ARM helix might be the missing structural element of the COMPASS eCM, which inhibits the enzymatic assembly upon binding to the nucleosome and is overcome by H2B~Ub for enzyme activation.

To validate this hypothesis, we first designed and prepared a COMPASS CM plus (CM+) complex by extending the Set1 SET domain with an extra ~30 amino acids at the N-terminus, which excludes the majority of the nSET domain except the arginine-rich motif. Strikingly, the activity of the COMPASS CM+ was similarly suppressed like the COMPASS eCM against naked nucleosome (Figure 7G). In a reaction against uNCPs as substrates, its activity was stimulated to an almost equal extent as the COMPASS eCM, indicating that the evolutionarily conserved Set1 ARM is indeed responsible for coupling H2Bub to enzyme activation (Figure 7G).

In company with the changes at the C-terminal half of the Set1 ARM helix, a profound difference between the uNCP- and NCP-bound COMPASS eCM structures can be found at the catalytic cleft of the Set1 SET domain. Despite the comparable resolution of the two structures, only three residues centered at the H3K4 methylation site show detectable density in the NCP-bound COMPASS structure, whereas all eight N-terminal residues of H3 can be clearly traced for the uNCP-bound form of the enzymatic complex (Figure 7H, S3C vs. S3D). By superimposing the nucleosomal portion of the two structures, we also detected a small, but considerable, degree of global structural changes within the Set1 catalytic domain, which narrow the catalytic cleft without impacting to the position of the SAM cofactor (Figure S7C, D). As a whole, the structural differences between the two structures might reflect the impact of H2B~Ub on the dynamic topology of the Set1 catalytic domain, which has been proposed to dictate enzymatic activity (Hsu et al., 2018; Li et al., 2016; Worden and Wolberger, 2019). We speculate that such an effect might be allosterically transduced through both the Ub-Swd1 and Ub-Set1 interfaces (Figure S7E).

### Conclusions

The cryo-EM structure of the COMPASS eCM bound to nucleosome reveals a proximity-enhanced substrate capture mechanism, in which the multi-subunit histone methyltransferase attaches itself to the nucleosome body to enable its catalytic subunit to catch the flexible N-terminal tail of H3 bearing the H3K4 methylation site. This substrate engagement mechanism is distinct from many other chromatin modifying enzymes, such as Ring1b-Bmi1-UbcH5c, SAGA-DUB, and DOT1L, which use the histone surface to precisely position their catalytic center in the immediate vicinity of their target sites (Figure S4) (Anderson et al., 2019; Jang et al., 2019; McGinty et al., 2014; Morgan et al., 2016; Valencia-Sánchez et al., 2019; Worden et al., 2019).

Binding of COMPASS to the nucleosome disk body not only serves the purpose of substrate recognition, but also renders the enzymatic complex susceptible for regulation. The COMPASS eCM-NCP complex is characterized by both extensive interactions among the subunits of the methyltransferase assembly and multiple interfaces made by four out of the six subunits with the nucleosomal disk. These contacts predict that local structural changes and spatial re-adjustments can propagate through the complex and affect the catalytic activity of the methyltransferase catalytic domain. Specifically, binding of COMPASS to the nucleosome traps the arginine-rich motif of Set1 at the interface, which adopts an α-helical conformation and docks to the nucleosomal acidic patch. By pushing against the Set1 catalytic domain sitting on top of histone H2A, the Set1 ARM helix might act as a pivot and inhibit COMPASS by affecting the spatial configuration of the Set1 catalytic domain relative to the rest of the COMPASS subunits. Upon conjugation to histone H2B, Ub stabilizes the C-terminal half of the Set1 ARM helix (Figure S7B) and packs against Swd1, which plays a key role in nucleating and organizing the COMPASS CM. These interactions conceivably enable Ub to override the inhibitory effect of the Set1 ARM helix by allosterically modulating the packing of the Set1 catalytic domain against Swd1, Bre2, and Swd3, thereby, altering the dynamic property of its active site.

Notably, in our enzymatic assay, the COMPASS eCM is more active than the full-length COMPASS complex towards both NCP and uNCP substrates (Figure 1B vs. 1D), suggesting that other components of the large N-terminal half of COMPASS that are missing in the COMPASS eCM also contribute to the suppression of enzymatic activity (Jeon et al., 2018; Kim et al., 2013). Furthermore, without the arginine-rich motif of the nSET domain, the COMPASS CM also has detectable response to H2Bub. It is plausible that H2Bub might facilitate the binding of the enzymatic assembly to nucleosomes by restricting the topological re-arrangement of COMPASS that favors nucleosome binding (Figure S8). Future studies will be needed to elucidate the detailed mechanisms of COMPASS regulation by H2Bub.

## Acknowledgements

We would like to thank Joel Quispe (UW Biochemistry EM facility) for assistance and training in EM sample preparation, microscope operation, and data collection. We would also like to thank the Colorado State University Protein Expression and Purification facility for individual histone proteins. The plasmids for expression of xenopus histones were a kind gift from Dr. Karolin Luger. This work was supported by the Howard Hughes Medical Institute (N.Z.), and the NIH (XXX to C.C.).

## Author contributions

P.L.H. and N.Z. conceived the project. P.L.H. purified all proteins, with the exception of individual histones, used in this study. H.S. screened initial cryo-EM grid conditions, and collected cryo-EM data with assistance from P.L.H. H.S. processed the cryo-EM data and generated maps for model building. Model building was conducted by P.L.H. with guidance from N.Z. P.L.H. conducted all binding and methyltransferase assays. C.L. and C.C. prepared large-scale nucleosomes used for structural studies. J.K. and C.C. prepared H2Bub-octamers for uNCP reconstitutions by P.L.H. Manuscript was written by P.L.H. and N.Z. with input from all authors.

## Declaration of Interests

The authors declare no competing financial interests. N.Z. is a member of the scientific advisory board of Kymera Therapeutics and a co-founder of Coho Therapeutics.

## METHODS

### Protein expression and purification

COMPASS subunits were PCR amplified from *K. lactis* genomic DNA and subcloned into modified pFastBac vectors for expression in insect cells. Recombinant viruses were produced as per manufacturer instructions, and subsequently amplified in Sf9 monolayer cells to generate high titer viruses for protein expression in HighFive monolayer cells. Purification of the COMPASS subomplexes and mutants were done as previously described (Hsu et al., 2018).

### Nucleosome reconstitution

Individual histones and the H3K4M mutant were expressed individually in BL21 DE3, and harvested 18 hours post-induction. Histone octamers were reconstituted using the “one-pot” refolding method as previously described (Lee et al., 2015). The 601-147 bp DNA sequence was excised from a plasmid containing 20 repeats of the sequence (McGinty et al., 2016). The plasmid backbone was removed using PEG precipitation, and the 147 insert was further purified by anion exchange chromatography, followed by desalting into TE buffer. Small scale titrations were done to determine a 1:1 DNA:histone ratio, followed up by a large scale dialysis to form nucleosomes (Dyer et al., 2004). Nucleosomes (WT and H3K4M) were stored at 4°C and used within a week of formation.

### H2Bub-nucleosome reconstitution

Individual xenopus histones, including the H3K4M and H2B K120C mutants, were purchased from the Protein Expression and Purification facility at Colorado State University (www.histonesource.com) as lyophilized powders. Ub G76C was expressed in BL21 DE3, and purified using a combination of acid precipitation followed by cation exchange chromatography. Purified Ub G76C was then dialyzed into water, and lyophilized for future use.

Generation of H2Bub was done essentially as previously described (Chatterjee et al., 2010). In brief, DTNP (2.0 mg, 6.45 mmol) was dissolved in 500 µL of a 3:1 (v/v) acetic acid:water mixture and added to H2B K120C (3 mg, 0.22 µmol). The reaction was allowed to proceed for 12 h at 25 °C, prior to purification by C4 semi-preparative RP-HPLC with a 30-70% B gradient. This yielded 2.1 mg (68.4%) of the pure H2B K120C-5-nitro-2-pyridinesulfenyl (pNpys) disulfide adduct. The identity of the disulfide adduct was verified by ESI-MS (expected: 13,622 Da, observed: 13,623 ± 3 Da). The product was then lyophilized for subsequent Ub ligation.

1 equivalent of the lyophilzed H2B-pNpys adduct (1.5 mg, 0.108 µmol) and 2.1 equivalents of Ub G76C (2.0 mg, 0.232 µmol) were dissolved in 625 µL of reaction buffer consisting of 1M HEPES, 6 M Gn-HCl, pH 6.9. Reaction was allowed for 1 h at 25°C with continuous shaking. The reaction products were purified by C4 semi-preparative HPLC with a 30-70% B gradient to yield H2Bub (1.2 mg, 74 %). The identity of the disulfide adduct was verified by ESI-MS (expected: 22,081 Da, observed: 22,083 ± 2 Da). The H2Bub product was then lyophilized and stored at −80°C for histone octamer reconstitution.

H2Bub-containing octamers were assembled using the salt dialysis method (Dyer et al., 2004) and purified over a Superdex 200 Increase 10/300 column (GE Healthcare) equilibrated in 10 mM HEPES pH 7.5, 2 M NaCl. Peak fractions were pooled and stored at −80°C for future use. H2Bub-nucleosomes were reconstituted as described for unmodified nucleosomes.

### COMPASS-nucleosome complex reconstitution

Purified COMPASS eCM subcomplex were added to 4-fold excess to NCP/uNCP, and allowed to mix at room temperature for 30 minutes before injection on to a Superdex 200 Increase 10/300 column (GE Healthcare) equilibrated in 10 mM HEPES pH 7.5, 100 mM NaCl, 1 mM DTT, 0.05 mM SAM (NEB). Peak fractions were collected, analyzed by SDS-PAGE, and pooled. The complex was concentrated to 5 mg/mL, flash frozen, and stored at −80°C for future use.

Preformed COMPASS eCM-nucleosome complexes were incubated with 0.1% (v/v) gluteraldehyde for 10 minutes at room temperature, and then quenched with the addition of 50 mM Tris pH 7.5. Crosslinked complexes were then injected on to a Superdex 200 Increase 10/300 (GE Healthcare) equilibrated in 20 mM Tris pH 7.5, 100 mM NaCl, 1 mM DTT, 0.05 mM SAM (NEB). Peak fractions were analyzed by negative stain electron microscopy, and the best fractions were pooled and concentrated to 0.6 mg/mL, and stored at −80°C for future grid preparation.

### Grid preparation and cryo-EM data collection

For cryoEM grid preparation, the complex (eCM-uNCP or eCM-NCP) was diluted to 0.3 mg/mL. 3 µL of the diluted material was applied to the holey side of a glow discharged (PELCO easiGlow) QuantiAu 1.2/1.3, 300 mesh grid (Quantifoil Micro Tools GmbH), and allowed to incubate for 1 minute at 4°C and 100% relative humidity. The grid was subsequently blotted for 3 seconds, plunge frozen into liquid ethane using a Vitrobot Mark IV system (Thermo Fisher Scientific), and stored in liquid nitrogen for data collection.

For eCM-uNCP, data collection was carried out on a Titan Krios transmission electron microscope (Thermo Fisher Scientific) operated at 300 kV, equipped with a post-column Gatan Quantum GIF energy filter and a Gatan K2 Summit direct detector. Movie stacks were recorded automatically using the Leginon software (Suloway et al., 2005) at a magnification of 130K, resulting a physical pixel size of 1.056 Å. The K2 camera was operated in super-resolution counting mode during data acquisition, and the slit width of energy filter was set to be 20 eV. A total dose of 74 e^−^/Å^2^ spanning a 9-second exposure time was fractionated into 45 frames. The whole dataset comprising 6,958 movie stacks was collected in two sessions with a defocus range set between 1 µm and 3 µm.

Settings of data collection for the eCM-NCP complex was similar to that used for eCM-uNCP. Either a 60-frame movie stack (3,855 movie stacks) or a 45-frame movie stack (10,336 movie stacks) was collected within a 9-second exposure, with a defocus range of 1 ~ 3 µm, and the total dose contained 74 e^−^/Å^2^. In total, 14,191 movie stacks were collected in five sessions.

### Cryo-EM data processing of COMPASS eCM-uNCP

All data processing steps were implemented within the RELION-3.0 pipeline (Kimanius et al., 2016; Zivanov et al., 2018) unless mentioned explicitly. Both dose-weighted summed images and non-dose-weighted summed images with a pixel size of 1.056 Å were generated through aligning movie frames using MotionCor2 (Zheng et al., 2017), and binned by a factor of 2 in both dimensions by Fourier cropping. Images were then imported into cisTEM (Grant et al., 2018) for manual inspection. Images with ice of bad qualities or of unreasonable thickness were excluded. A subset of 6,036 high quality images was used for automatic particle picking in cisTEM, resulting in a set of 426,260 particles. The particle coordinates were exported in the Relion format for further processing. Parameters of the contrast transfer function (CTF) were estimated on the non-dose-weighted summed images by GCTF (Zhang, 2016). Particles were then extracted from the dose-weighted summed images.

A stack of 357,811 particles was created after data cleaning by 2 rounds of reference-free 2D classification. Global 3D classification without masking provided an initial model generated from RELION-3.0 was then performed to further classify the particles into 4 classes, in which the uNCP and one copy of the COMPASS eCM were better resolved, while the other copy of COMPASS eCM was less clear. To obtain a better class of eCM-uNCP complex, the set of 357,811 particles after 2D classification was subjected to a focused 3D classification procedure with a soft mask imposed, which enclosed both the uNCP and the better resolved copy of COMPASS eCM. Particles belonging to the best class from the last 6 iterations were combined, and duplicated particles were removed, giving rise to a particle set of 215,199 particles. This subset of particles went through focused 3D refinement with the same mask applied, resulting in a 3.6 Å density map. To further improve the quality of the reconstruction, particles used in the focused 3D refinement were imported for beam tilt parameters estimation in the CTF refinement job type (Zivanov et al., 2018) in RELION-3.0. The final density map was refined to a resolution of 3.5 Å by another round of focused refinement. The post-processing protocol in RELION-3.0 was used to correct for the modulation transfer function, sharpen the density map, and calculate the gold-standard Fourier shell correlation (FSC). According to the FSC 0.143 criterion (Rosenthal and Henderson, 2003), the reported resolution of the map is 3.5 Å. The map was sharpened by applying a global B-factor of −123 Å^2^ estimated during post-processing. Local resolution was determined by ResMap (Kucukelbir et al., 2014).

### Cryo-EM data processing of COMPASS eCM-NCP

Image processing of COMPASS eCM-NCP was similar as that of eCM-uNCP. Movie frame alignment was carried out using MotionCor2 (Zheng et al., 2017), and binned by a factor of 2 in both dimensions by Fourier cropping. Both dose-weighted summed images and non-dose-weighted summed images with a pixel size of 1.056 Å were generated for later process. Summed images were imported into cisTEM (Grant et al., 2018) for manual inspection. A subset of 12,342 images with good ice quality and thickness were selected for later processing. 1,547,866 particles were automatically picked in cisTEM (Grant et al., 2018) on the good images, and coordinates were exported to RELION-3.0 for particle extraction. CTF parameters estimation for each individually picked particle were performed by GCTF (Zhang, 2016).

A pool of 1,547,866 particles were extracted from dose-weighted summed images and subjected to 2 rounds of reference-free 2D classification for particle clean up. A subset of 955,956 particles was obtained by selecting good 2D class averages. Initial model was created in RELION-3.0 and used for global 3D classification without masking. After merging and removing duplicated of particle assigned to the best class from the last 3 iterations, 221,787 particles were refined with a soft mask applied, which encompassed both the nucleosome core particle and a copy of the COMPASS eCM complex that showed a clearer density. A 3.76 Å density map was produced from the 3D refinement. To obtain a better density map, CTF refinement (Zivanov et al., 2018) was employed to estimate the beam tilt parameters, similar to that was done for the eCM-uNCP complex. In addition, Bayesian polishing (Zivanov et al., 2019) in RELION-3.0 was carried out to generate shiny particles. The set of 221,787 shiny particles were further processed by a second round of focused global 3D classification, with a finer angular search step (3.7°) and a τ value of 20. The final dataset comprised of 100,905 shiny particles and a 3.7 Å density map was generated after 3D auto-refine. The reconstructed map was post-processed in RELION-3.0. A global B-factor of −91 Å^2^ estimated during post-processing was applied to sharpen the map. The final resolution was determined to be 3.7 Å based on the FSC 0.143 criterion (Rosenthal and Henderson, 2003). ResMap (Kucukelbir et al., 2014) was used for local resolution determination.

### Model building and refinement

PDB crystal structures for the COMPASS CM (PDB: 6CHG), Ubiquitin (PDB: 1UBQ), NCP assembled with a 601 positioning sequence (PDB: 3MVD), and a homology model for *K.lactis* Spp1 generated from an HHPred alignment (Söding et al., 2005) and Modeller (Eswar et al., 2006) were rigid-body fit into the COMPASS eCM-uNCP density map using Chimera (Pettersen et al., 2004). The models were then manually trimmed, extended, and adjusted in Coot (Emsley and Cowtan, 2004). Models were then subject to several rounds of both local and global real-space refinement in Coot and Phenix (Adams et al., 2010). Statistics of the final refinement are reported in Table S1. Model building and refinement were done similarly for the COMPASS eCM-NCP complex.

### Methyltransferase assays

1 µM of enzyme and 0.5 µM of nucleosome/H2Bub-nucleosome substrates were incubated in buffer containing 20 mM HEPES pH 7.5, 100 mM NaCl, and 0.2 mM SAM (NEB) for 15 minutes at 30°C. Reactions were quenched with the addition of SDS-PAGE loading buffer, resolved on a 15% gel, and transferred to PVDF membranes. Membranes were blocked at room temperature in 5% milk in TBS-T, and then probed with antibodies against H3 (Abcam ab1791), H3K4me1 (CST D1A9), H3K4me2 (Abcam ab7766), and H3K4me3 (Abcam ab8580) overnight at 4°C. Blots were washed the next morning with TBS-T and then probed with secondary antibody at room temperature for 1 hour. Blots were subsequently washed with TBS-T to remove excess secondary. The blots were then incubated with ECL reagent, and exposed on an imager for analysis.

### Nucleosome binding assays

50 nM H3K4M-nucleosome/H2Bub-nucleosome were incubated with increasing concentrations of COMPASS complexes (0.1 µM, 0.25 µM, 0.5 µM, 1.0 µM, 2.5 µM, 5.0 µM) in a buffer containing 20 mM HEPES pH 7.5, 50 mM NaCl, 0.05 mM SAM (NEB) for 30 minutes at room temperature. Glycerol was then added to a final concentration of 10%, and then loaded into a pre-chilled, pre-run 5% (29:1) 0.2X TBE native acrylamide gel. Gels were run at 120V for 90 minutes, and then stained with SybrGold, and exposed on an imager for analysis.

### Quantification and statistical analysis

Protein quantification was done using Bio-Rad Protein Assay Dye, and comparing the readings against a standard curve generated using BSA. Nucleosome concentrations were determined using an A260 extinction coefficient of 2,784,500 M^−1^ cm^−1^ on a Nanodrop spectrophotometer (Thermo Fisher).

### Data Availability

The accession number for the cryo-EM map of the COMPASS eCM-uNCP, and eCM-NCP complex are EMDB-XXXX and EMDB-XXXX, respectively. The coordinates for the COMPASS eCM-uNCP, and eCM-NCP is deposited under the accession numbers PDB: XXXX and PDB:XXXX, respectively.

## Supplemental Figure Titles and Legends

**Figure S1.**
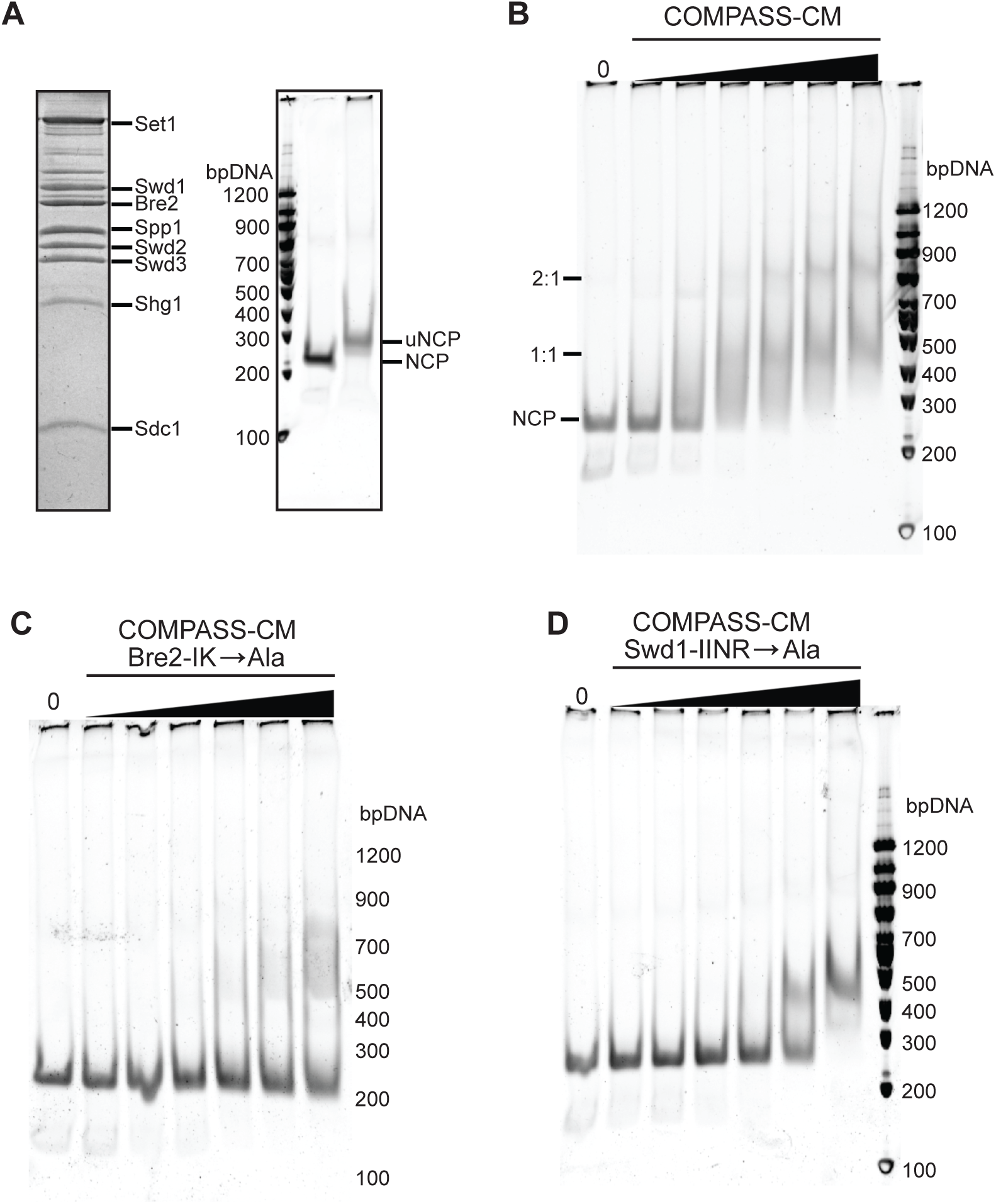
Nucleosomal binding by COMPASS and its subcomplexes (related to Figure 1, Figure 3, Figure 4) A. Recombinant *K. lactis* COMPASS purified from insect cells are resolved on SDS-PAGE and stained with Coomassie. All eight subunits are used in their full-length forms with the exception of Sdc1, which comprises only the dimerization domain (left panel). Unmodified and H2Bub nucleosomes (NCP and uNCP) run on a native TBE gel, and stained with SybrGold (right panel).
B. Native TBE gel shift assay of unmodified nucleosomes with increasing concentrations of COMPASS-CM, stained by SybrGold.
C. Native TBE gel shift assay of unmodified nucleosomes with increasing concentrations of COMPASS-CM assembled with a Bre2-IK→AA mutant, stained by SybrGold.
D. Native TBE gel shift assay of unmodified nucleosomes with increasing concentrations of COMPASS-CM assembled with a Swd1-IINR→A4 mutant, stained by SybrGold.

**Figure S2.**
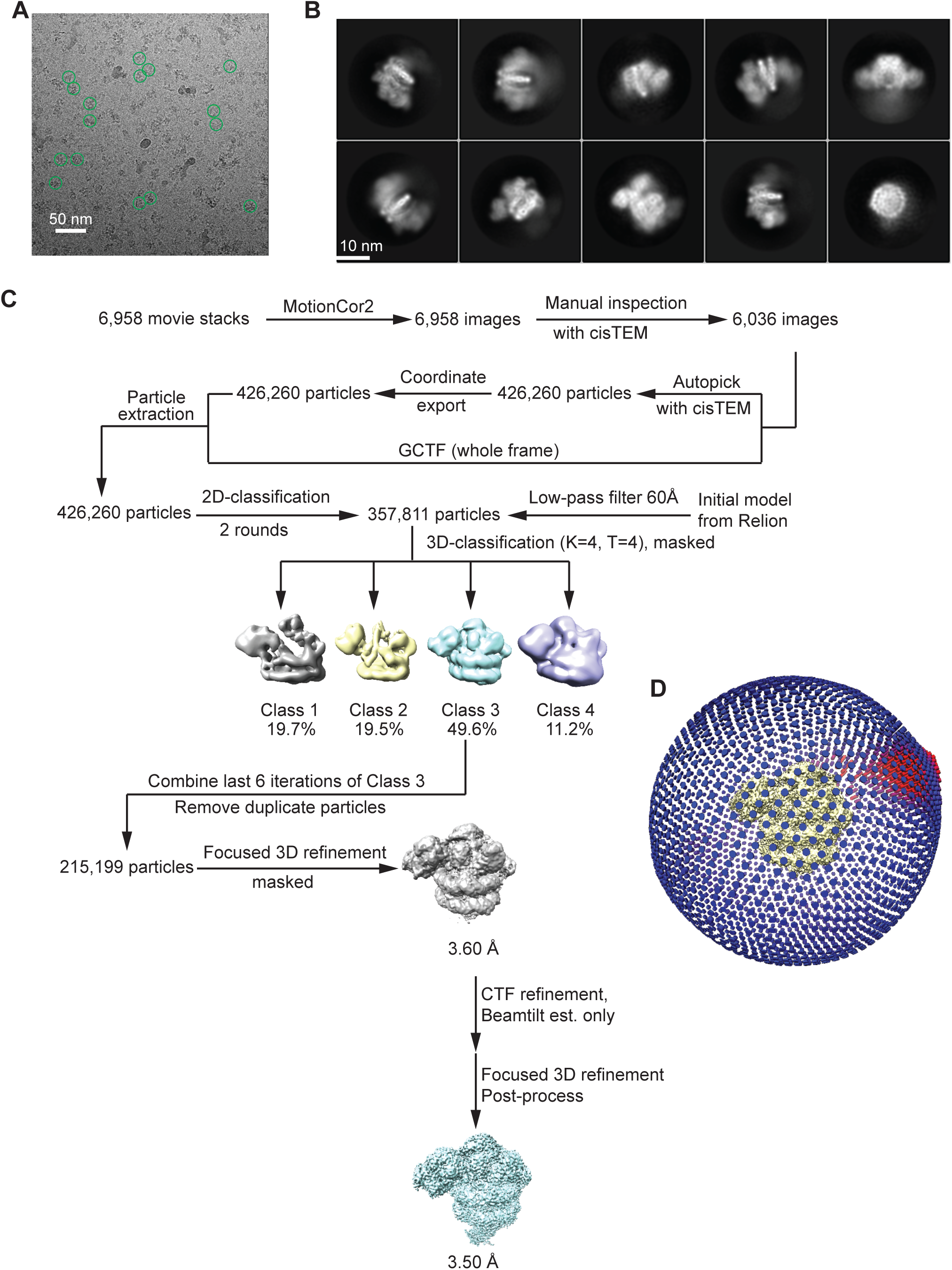
Cryo-EM analysis of the COMPASS eCM-uNCP complex (related to Figure 2) A. A representative cryo-EM image of the COMPASS eCM-uNCP complex with typically picked particles circled in green. Scale bar, 50 nm.
B. Representative 2D class averages of the cryo-EM dataset, revealing different orientations of the particles. Two copies of the COMPASS eCM bind to one copy of the H2Bub nucleosome with one copy of the eCM displaying weaker density. Scale bar, 10 nm.
C. Data processing flow chart for the COMPASS eCM-uNCP complex.
D. Angular distribution of the particles used in the final reconstruction. The density map is shown in yellow in the center of the distribution.

**Figure S3.**
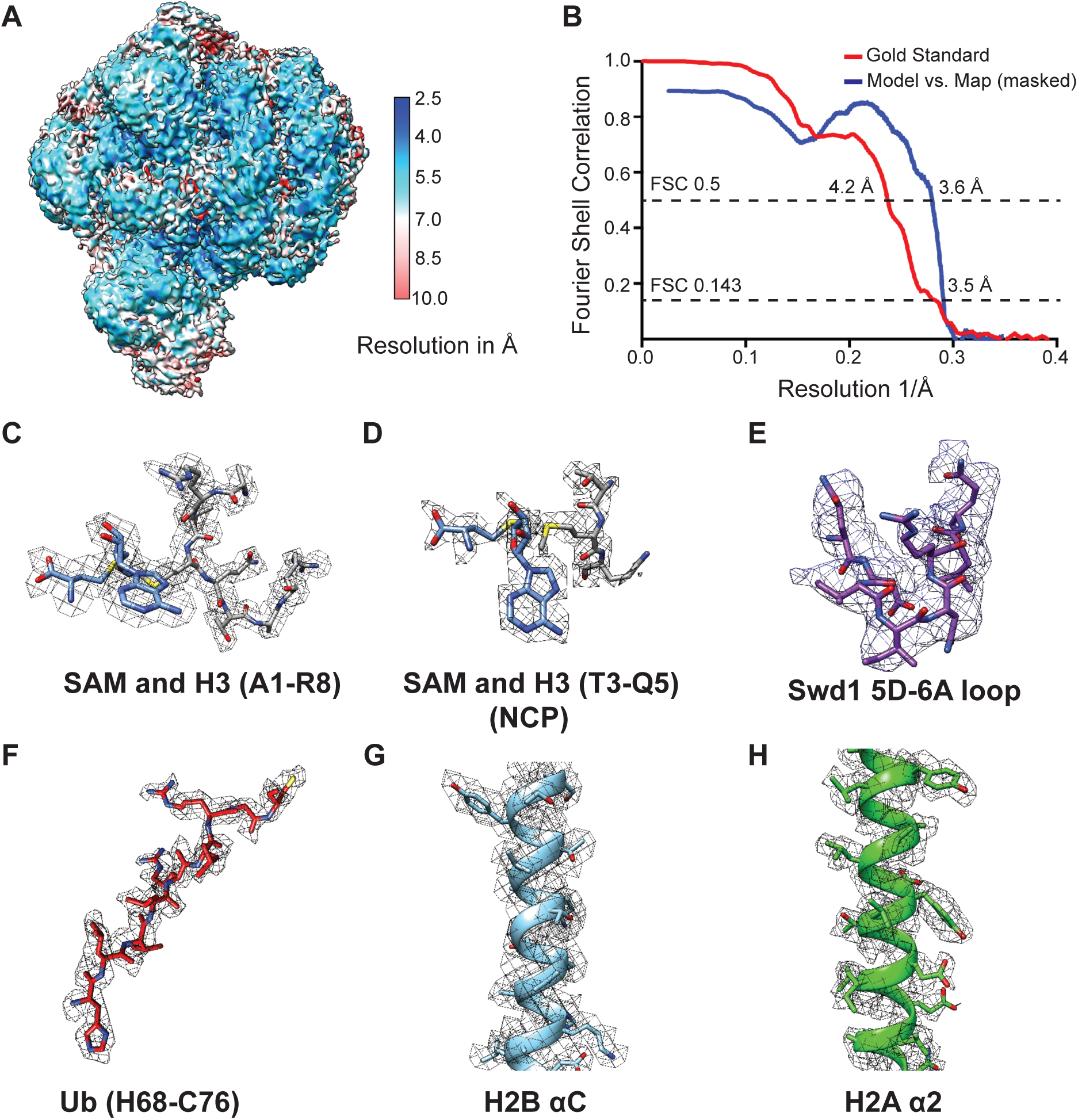
Structure model quality of the COMPASS eCM-uNCP complex (related to Figure 2, Figure 7) (A) Local resolution assessment by ResMap for the COMPASS eCM-uNCP complex. The local resolution in Å is displayed on the final masked map that contains only a 1:1 complex.
(B) Gold-standard and model vs. map FSC curves for both the final reconstruction and final refinement. The final map resolution is 3.5 Å as calculated by the FSC 0.143 standard. The final model resolution is estimated to be 3.6 Å as judged by a FSC 0.5 standard.
(C) Density of the SAM cofactor and H3 tail peptide found in the Set1 active site of the COMPASS eCM-uNCP structure.
(D) Density of the SAM cofactor and H3 tail peptide found in the Set1 active site of the COMPASS eCM-NCP structure.
(E) – (H) Representative densities of key structural elements found in the COMPASS eCM-uNCP structure.

**Figure S4.**
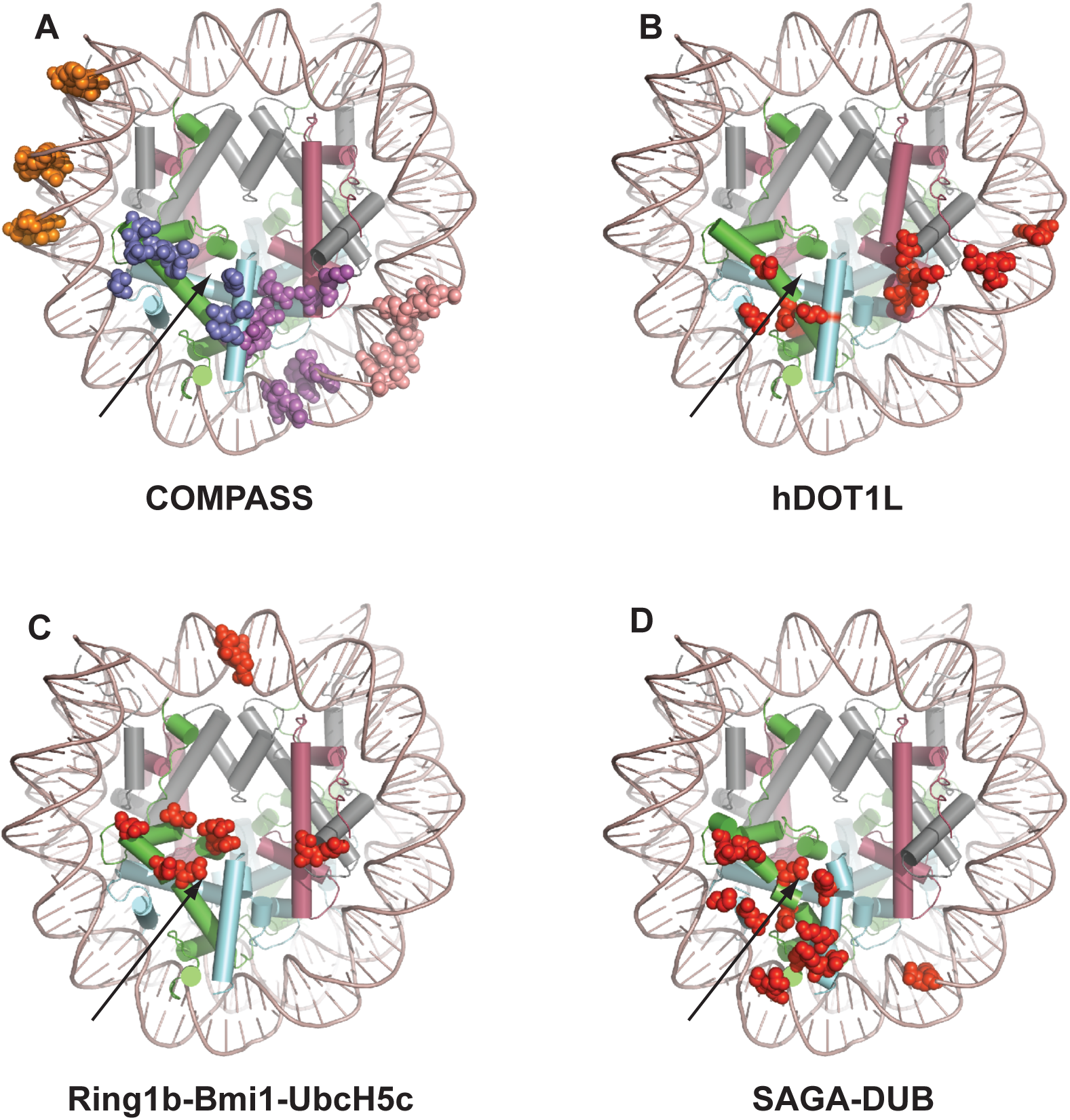
Nucleosome disk face binding mode comparisons (related to Figure 2) (A) View of the nucleosome looking down the disk face. H3 is in dark grey, H4 in deep red, H2A in green, and H2B in cyan. Contact points between the nucleosome and the COMPASS eCM are shown in spheres. Spheres are colored according to the same subunit coloring scheme as in Figure 2. The acidic patch position is indicated by an arrow.
(B) – (D) View of nucleosome contacts between several representative chromatin binding proteins/enzymes. Points of interactions are colored as red spheres. Histone coloring scheme is the same as in (A). The acidic patch position is indicated by an arrow.

**Figure S5.**
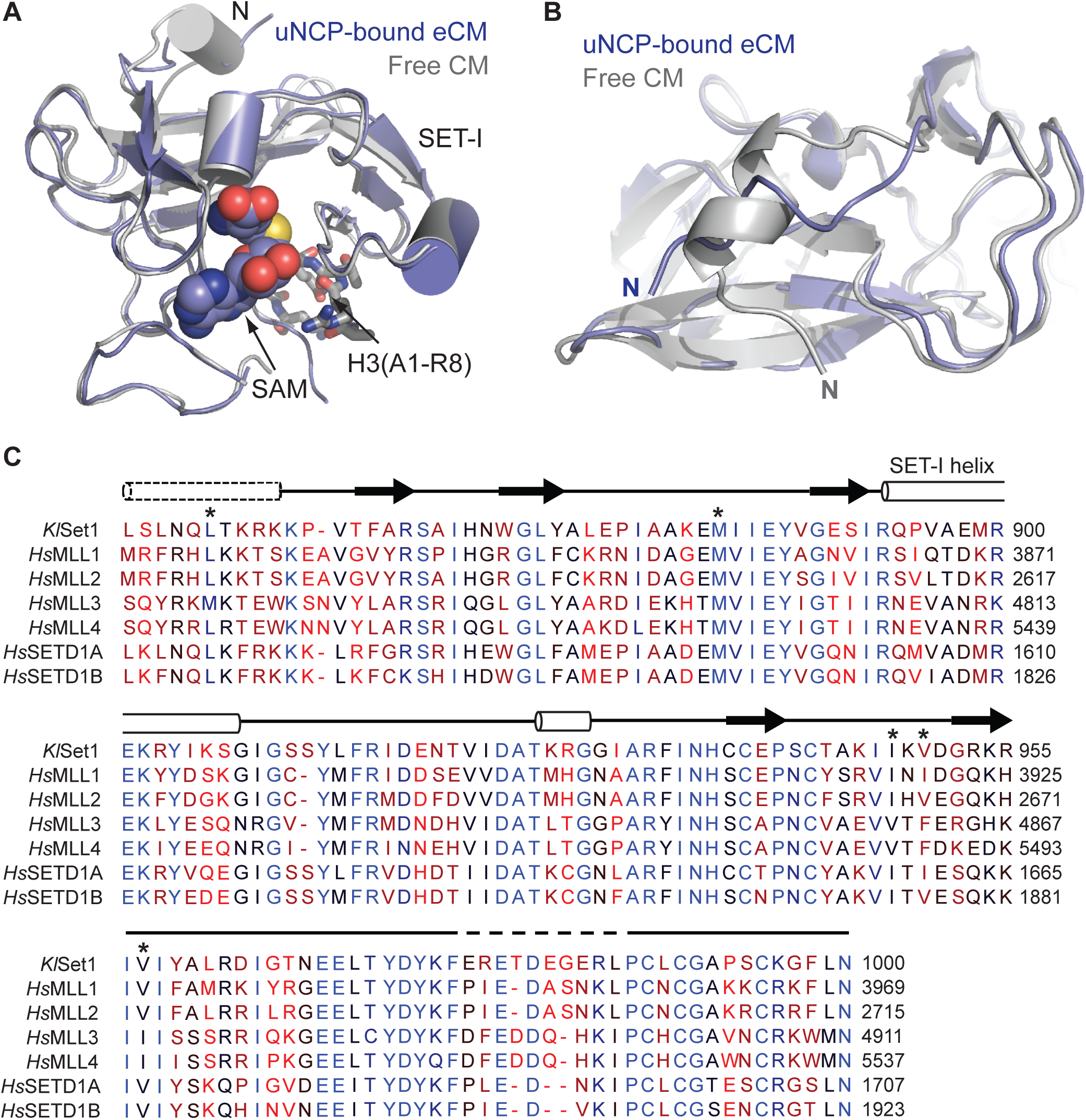
Conserved structural features of Set1 (related to Figure 5) A. Superposition of the SET domain of Set1 in the context of an uNCP-bound eCM (blue) and free CM (PDB: 6CHG) (grey). SAM is shown in space filling spheres, and the H3 peptide is shown in grey sticks.
B. Superposition of the SET domain of Set1 in the free (grey, PDB: 6CHG) and NCP bound (blue) state, highlighting the conformation of the SET N-terminal helix. The position of the extreme N-terminus is labeled.
C. Structure-based alignment of yeast Set1 and human SET1/MLL SET domains. Cylinders denote helices, arrows show β-strands, and lines are loops. Dashed lines are regions not modeled in the structure. The dashed helix is the structural element that undergoes a conformational change upon nucleosome binding. Residues with asterisks above form the hydrophobic claw used to interact with H2A. The mobile SET-I helix involved in enzymatic activity is labeled.

**Figure S6.**
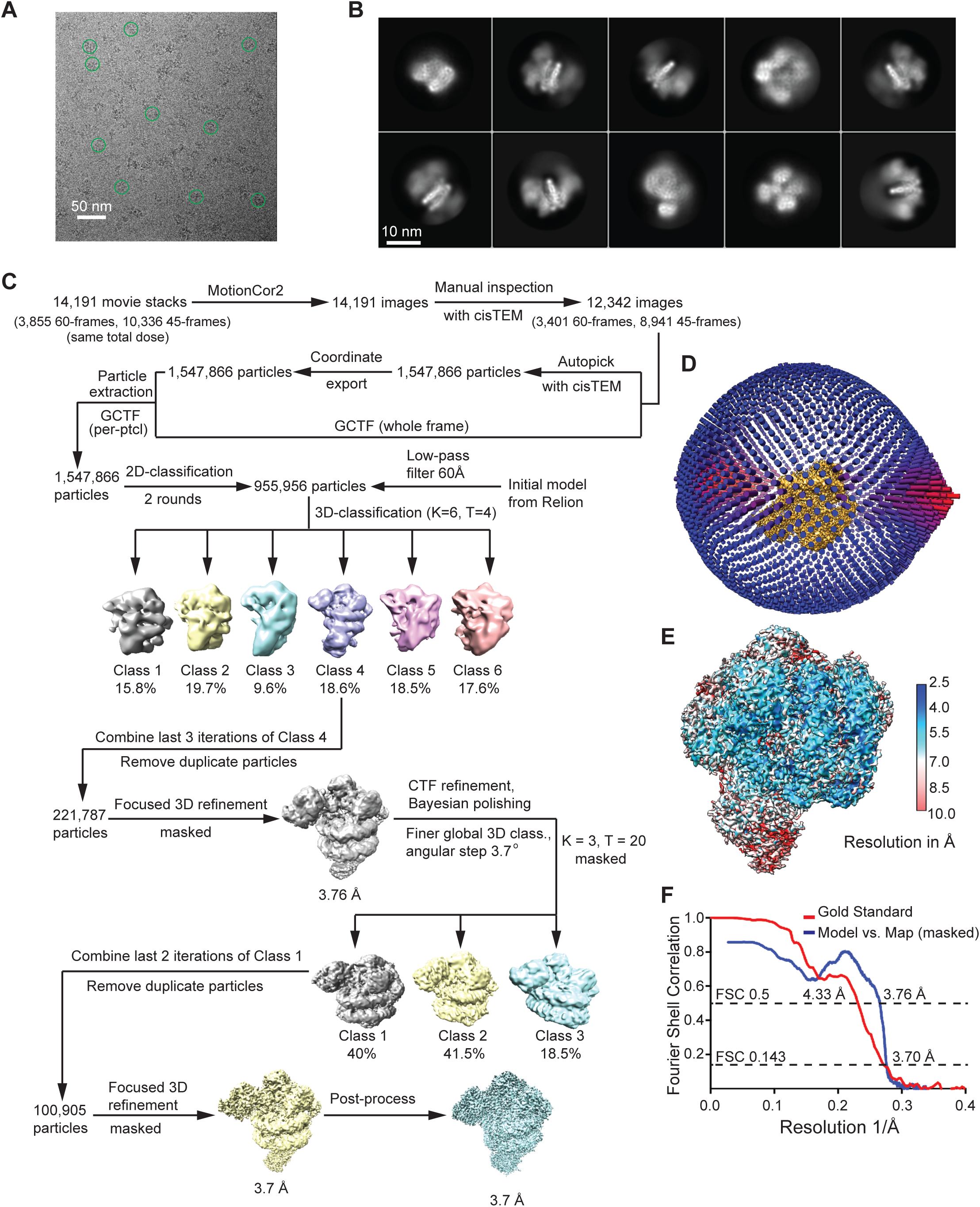
Cryo-EM analysis of the COMPASS eCM-NCP complex (related to Figure 7) A. A representative cryo-EM image of the COMPASS eCM-NCP complex with typically picked particles circled in green. Scale bar, 50 nm.
B. Representative 2D class averages of the cryo-EM dataset, revealing different orientations of the particles. Two copies of the COMPASS eCM bind to one copy of the nucleosome, with one copy of the eCM displaying weaker density. Scale bar, 10 nm.
C. Data processing flow chart for the COMPASS eCM-NCP complex.
D. Angular distribution of the particles used in the final reconstruction. The density map is shown in yellow in the center of the distribution.
E. Local resolution assessment by ResMap for the COMPASS eCM-NCP complex. The local resolution in Å is displayed on the final masked map that contains only a 1:1 complex.
F. Gold-standard and model vs. map FSC curves for both the final reconstruction and final refinement. The final map resolution is 3.7 Å as calculated by the FSC 0.143 standard. The final model resolution is estimated to be 3.76 Å as judged by a FSC 0.5 standard.

**Figure S7.**
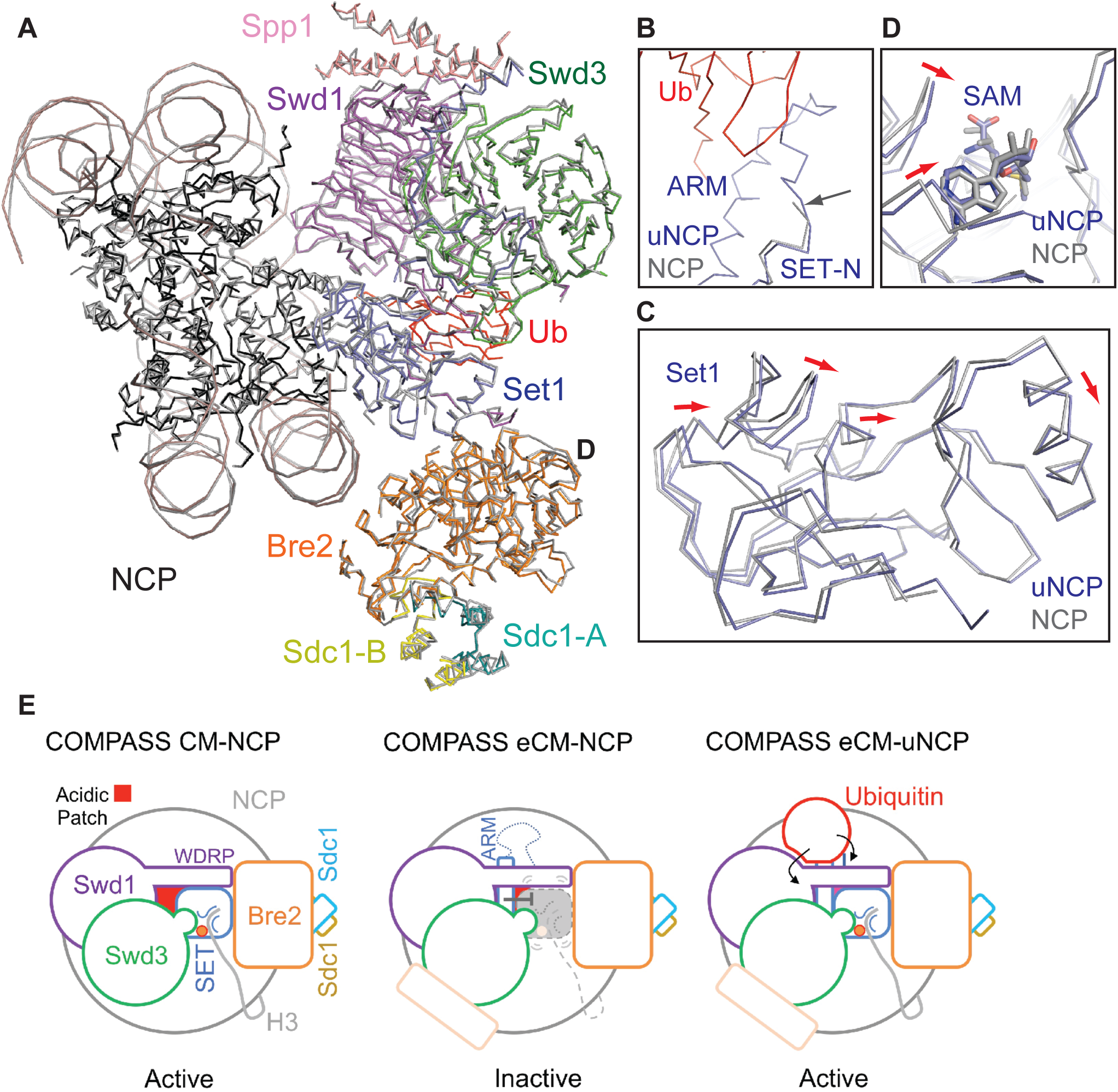
Structural comparison of the COMPASS eCM bound to uNCP/NCP (related to Figure 7) A. Superposition of the COMPASS eCM bound to H2Bub-(colored as in Figure 2) and unmodified-nucleosomes (eCM in grey, histones in black) via superimposing the nucleosome. All subunits shown in ribbon representation. Histones superposition near perfectly.
B. Close up view of the junction between the Set1 (blue) ARM, SET-N, and Ub (red). An arrow indicates where the N-terminus of the SET domain in the NCP structure ends, illustrating how Ub stabilizes the conformation of the loop and ARM.
C. Superposition of the Set1 SET domain, looking down the active site. Red arrows indicate the overall shift of the backbone upon the engagement of an H2Bub nucleosome.
D. Superposition of the SAM cofactor between the uNCP (blue) and NCP (grey) structures. Red arrows indicate shifts in the backbone after binding to an H2Bub nucleosome. Note the position of SAM is relatively unchanged between the two structures.
E. A schematic model of H2Bub-dependent H3K4 methylation by COMPASS. In the context of the CM, which is shared among all SET1/MLL enzymes, COMPASS is capable of methylating H3K4 in the absence of Ub. In the context of COMPASS specific eCM, the Set1 ARM helix is wedged between the nucleosomal acidic patch and COMPASS, and inhibits the activity of the SET domain, potentially through local structural perturbation. In the presence of H2B~Ub, Ub stabilizes the ARM helix and packs against Swd1. These local interactions are transduced to the Set1 SET domain, alleviating its inhibition by the ARM helix.

**Figure S8.**
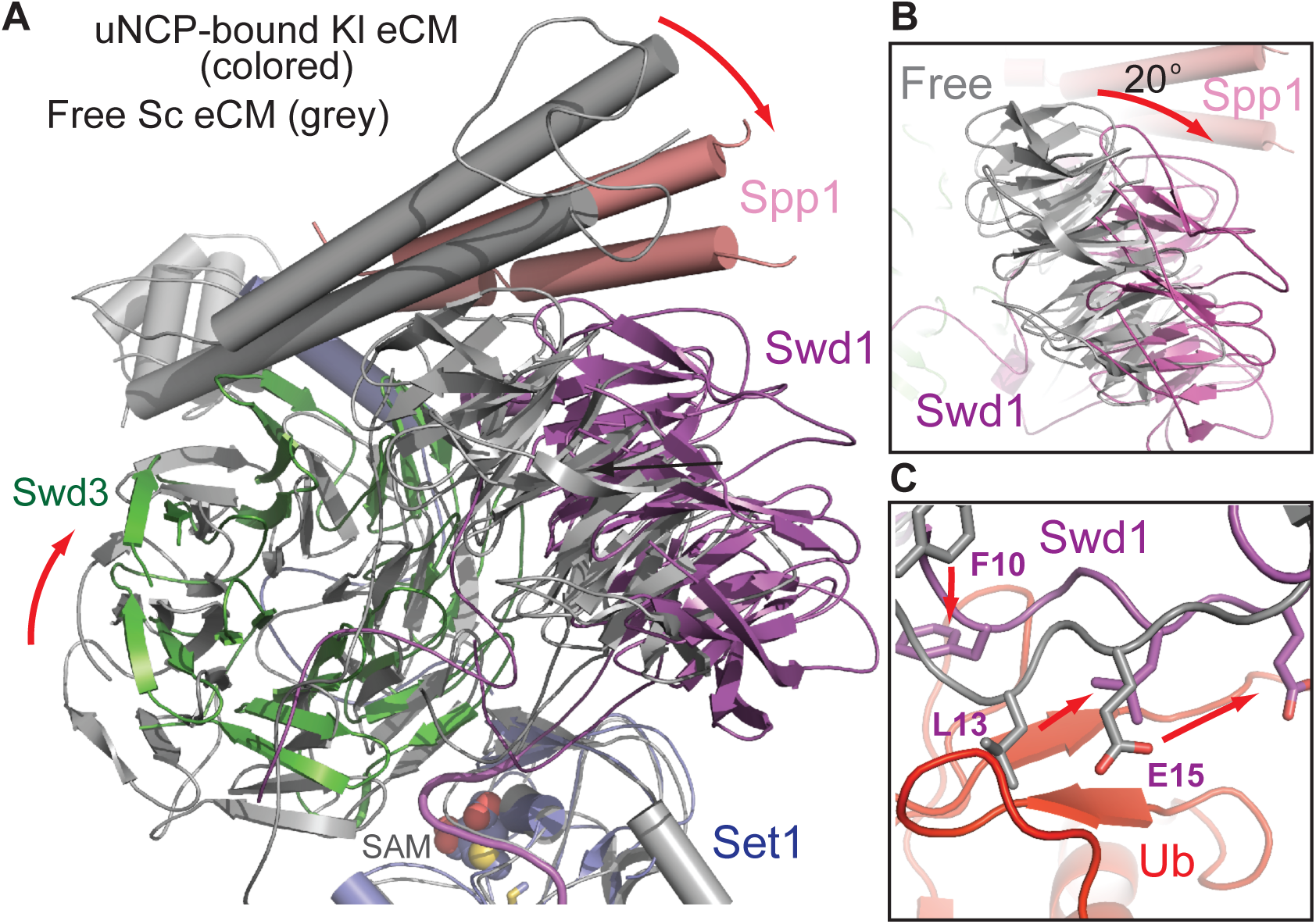
A topology of free COMPASS is incompatible with Ub binding. A. Superposition of the uNCP-bound *K. lactis* eCM (colored as in Figure 2) and free *S. cerevisiae* eCM (PDB: 6BX3) (grey) aligned on the SET domain. SAM is shown in spheres. Red arrows indicate the shift in conformation upon uNCP binding.
B. Close up view of the Swd1 WD40 domain between the free (grey) and uNCP bound (Swd1: magenta, Swd3: green, Spp1: pink) forms of the COMPASS eCM. Note the rotation of the propeller between the two different states.
C. Close up view of the Swd1 N-terminus between the free (grey) and uNCP bound (magenta) forms of the COMPASS eCM. Ub is shown in red. Phe10, Leu13, and Glu15 (numbered according to *K. lactis* ortholog) are shown in sticks. Note the clash of Leu13 against Ub in the free form of *S. cerevisiae* COMPASS eCM previously captured (Qu et al., 2018). Arrows note the shift of these residues upon nucleosomal binding.

**Table S1.**
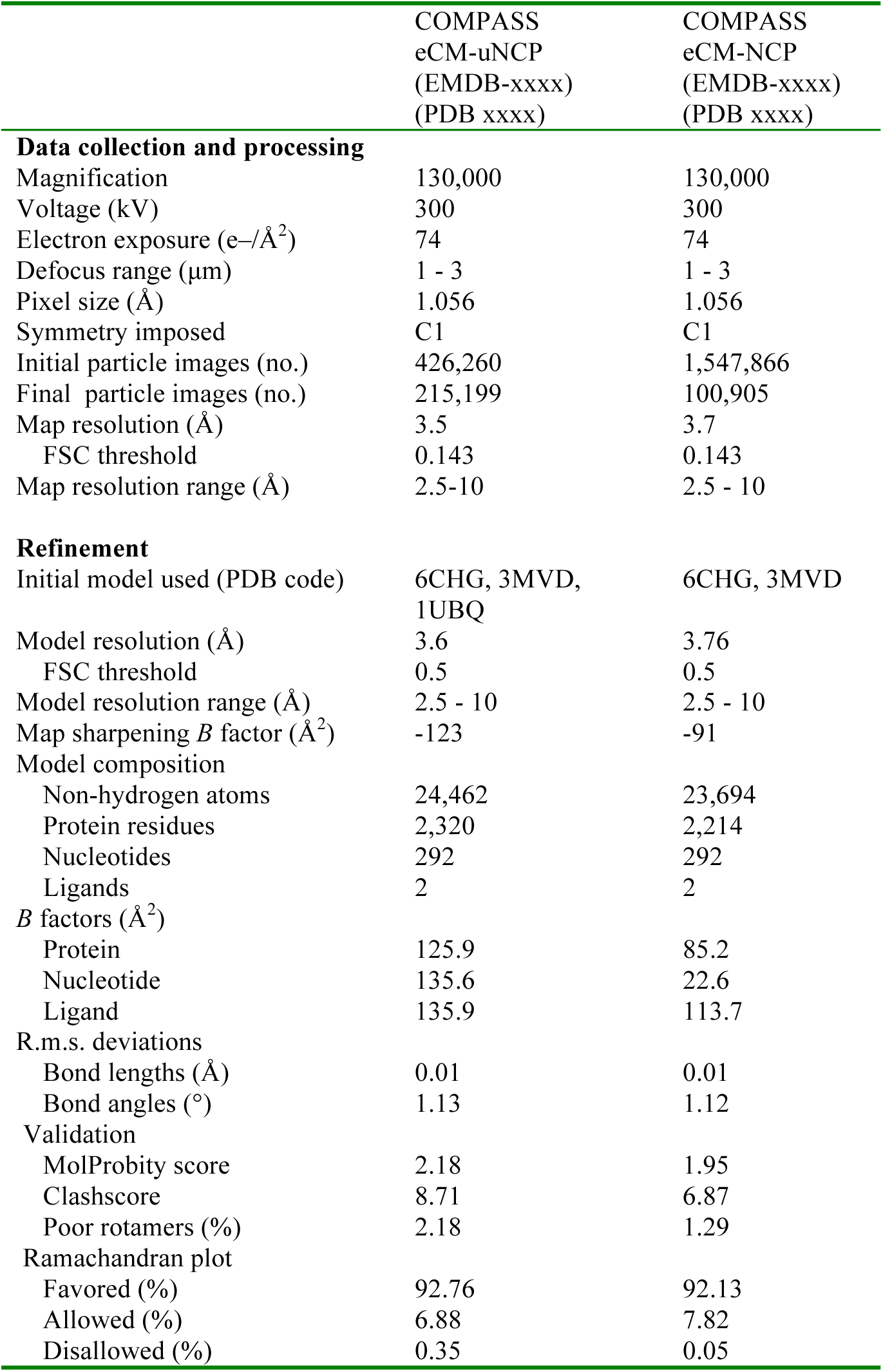
Cryo-EM data collection and model refinement statistics (related to Figure 2)

